# *LORELEI* sequence variation modulates lineage-dependent pollen tube reception in Brassicaceae

**DOI:** 10.64898/2026.11.18.767685

**Authors:** Emma Jong, Jennifer A. Noble, Matthew J. Alcaraz, Iz Varghese, Said Hafidh, David Honys, George P. Lisi, Mark A. Beilstein, Ravishankar Palanivelu

**Affiliations:** The School of Plant Sciences, University of Arizona, Tucson, AZ 85721, USA; Institute of Experimental Botany of the Czech Academy of Sciences, Rozvojová 263, 165 02 Prague 6, Czech Republic; Department of Molecular Biology, Cell Biology, and Biochemistry, Brown University, Providence, RI 02903, USA

**Keywords:** pollen tube reception, prezygotic reproductive barrier, LORELEI (LRE), Brassicaceae, reproductive isolation

## Abstract

Pollen tube reception (PTR) is essential for fertilization and can act as a prezygotic barrier in interspecific crosses. While core female components such as LORELEI (LRE) and FERONIA (FER) mediate conspecific PTR, their roles in heterospecific compatibility remain unclear. Here, we investigated how LRE sequence variation influences PTR outcomes with heterospecific pollen in Brassicaceae. In *Arabidopsis thaliana*, PTR frequently failed when ovules were pollinated with *Arabidopsis lyrata* or *Sisymbrium irio*, and loss of LRE or FER further increased pollen tube overgrowth in a pollen-species-dependent manner. Complementation of *lre* ovules with LRE orthologs showed distinct, donor-dependent effects: *AlyLRE* and *SiLRE* produced modest and variable improvements with *A. lyrata* pollen, whereas SiLRE consistently and markedly improved reception of *S. irio* pollen. Evolutionary analyses indicated relaxed purifying selection in the LRE lineage relative to its paralog *LLG1* and nominated residues 47 and 120 as candidates for functional divergence. Consistent with this, the Lineage II-associated R120T substitution enhanced reception of *S. irio* pollen while partially reducing conspecific PTR. In addition, the ancestral-state E47P–R120G combination also improved heterospecific PTR without measurably compromising conspecific function. Together, these results indicate that LRE sequence diversification can modulate female-side compatibility with pollen tubes from different evolutionary lineages, potentially by altering how pollen tube-derived signals are sensed and transduced during reception.

## Introduction

Reproductive isolation results from both prezygotic and postzygotic barriers that prevent successful hybridization between species. Prezygotic barriers act before fertilization, either via disrupting pollen-pistil interactions or by preventing zygote formation, while postzygotic barriers act after fertilization by affecting hybrid seed viability (Baack et al. 2015). In angiosperms, prezygotic barriers can occur before pollination, because of traits such as flower color and pollinator preference, or after pollination, during processes such as pollen hydration, germination, pollen tube growth, guidance, and reception in the pistil. Understanding the molecular mechanisms underlying these post-pollination barriers is important for explaining how species-specific reproductive compatibility is achieved and for informing breeding programs designed to generate novel hybrids.

When fertilization occurs, the pollen grain germinates on the stigma and forms a pollen tube, which then grows through the style and transmitting tract to reach an ovule. Once the pollen tube enters the micropyle, it contacts a synergid cell, which halts its growth and triggers sperm release in a process called pollen tube reception (PTR). Each step in this pathway, from pollen germination to PTR, can serve as a prezygotic barrier to fertilization. Studies of pollen-pistil interactions within a species (conspecific interactions) have identified several molecular components necessary for effective fertilization, and variation in these components can result in reproductive incompatibility. Consequently, molecular mechanisms that facilitate compatible pollen-pistil interactions may also act as barriers when pollen and pistils originate from different species (heterospecific interactions).

In *Arabidopsis thaliana*, pollen hydration and germination during compatible pollination are regulated by a receptor signaling module that includes the CrRLK1L receptor kinases FERONIA (FER) and ANJEA (ANJ), the co-receptor LORELEI-LIKE GPI-ANCHORED PROTEIN 1 (LLG1), and peptide ligands. Before pollination, stigma-derived RAPID ALKALINIZATION FACTOR 23 and 33 (RALF23 and RALF33) peptide ligands maintain FER–ANJ-dependent production of reactive oxygen species (ROS) in papilla cells. During compatible pollination, pollen coat-derived POLLEN COAT PROTEIN B-class peptides (PCP-Bs) compete with RALF23 and RALF33 for binding to the receptor complex, reducing stigmatic ROS and promoting pollen hydration and germination (Liu et al. 2021). This ligand switch also promotes water availability to the pollen by activating plasma-membrane H+-ATPases and opening PIP aquaporin channels (Liu et al. 2025). FER-dependent ROS signaling is also recruited during self-incompatibility in *Brassica rapa*, where self-pollination increases stigmatic ROS, whereas compatible pollination reduces it. Disruption of FER–Rac/Rop signaling, RBOH activity, or ROS accumulation compromises self-pollen rejection (Zhang et al. 2021). Subsequent work linked this pathway to the classical self-incompatibility recognition system by showing that binding of the pollen ligand SCR/SP11 to SRK promotes SRK association with FER, thereby linking self-pollen recognition to FER-dependent ROS signaling (Huang et al. 2023).

Related signaling mechanisms add to the rejection of pollen from other species. In *A. thaliana*, stigma-derived RALFs act through CrRLK1L receptors and LRX proteins to maintain a barrier to pollen tube penetration, while RALFs from compatible pollen compete with these stigma-derived ligands and weaken the barrier. Disrupting this pathway, or applying synthetic pollen-derived RALFs, can allow otherwise incompatible pollen tubes to penetrate the stigma and, in some cases, result in hybrid embryo formation (Lan et al. 2023). Additional mechanisms reinforce this interspecific barrier: the stigma-localized transmembrane protein STIGMATIC PRIVACY 1 (SPRI1) restricts penetration by pollen from distantly related Brassicaceae species independently of the self-incompatibility pathway (Fujii et al. 2019), and *SPRI2/SRS7* and its paralog *SPRI2*-like/*SRS5* further strengthen this barrier by regulating SPRI1 and the cell-wall-associated genes *TBL40* and *TBL45* (Fujii et al. 2023). Recently, pollen-derived SRK-interacting interspecific pollen signals (SIPS) were shown to activate an interspecific stigma barrier by engaging SRK, promoting its association with FER, and stimulating FER-dependent ROS production; loss of these pollen signals weakens rejection of heterospecific pollen (Cao et al. 2025). Thus, signaling systems that regulate compatible pollen acceptance can be differentially engaged during self-incompatible and heterospecific interactions, converting components of normal pollen-stigma communication into checkpoints for reproductive compatibility (Zhang et al. 2026).

After germination, the pollen tube must traverse the stigma and grow through the pistil’s transmitting tissues to reach the ovary, providing additional opportunities to reject pollen from other species. Researchers have identified several molecular components of this stylar barrier, particularly in *Solanum*. Pistil-expressed S-RNases and High Top-band (HT) proteins contribute to heterospecific pollen rejection, while pollen-side factors such as S-locus F-box proteins and CULLIN1 can confer resistance to these barriers. Suppression of HT proteins can weaken or eliminate stylar rejection of heterospecific pollen, showing the contribution of multiple female factors to this barrier (Tovar-Mendez et al. 2017). Other extracellular components, including stigma/style cysteine-rich adhesins and related lipid-transfer proteins, facilitate pollen tube adhesion and growth through the transmitting tract and may likewise influence species-dependent compatibility (Wang and Filatov 2023). Thus, both female stylar factors and pollen-expressed components determine whether a heterospecific pollen tube can successfully navigate the pistil, making pollen–style interactions an additional molecular checkpoint in post-pollination reproductive isolation.

An additional critical post-pollination checkpoint is pollen tube guidance to the ovule. Peptides produced by the female gametophyte attract conspecific pollen tubes from the transmitting tract toward the ovular micropyle, and some of these guidance signals display species-specific preferences. For example, in maize, the synergid-expressed peptide EGG APPARATUS 1 (ZmEA1) attracts pollen tubes in a species-preferential manner; when ZmEA1 is expressed in *A. thaliana* ovules, it can direct maize pollen tubes toward the micropyle (Márton et al. 2012). Similarly, in Torenia and *A. thaliana*, synergid-secreted LURE peptides preferentially attract pollen tubes from the same species (Higashiyama et al. 2006; Takeuchi and Higashiyama 2012). In *A. thaliana*, the receptor kinase PRK6, located at the pollen tube tip, detects AtLURE1 and is required for efficient LURE1-mediated guidance (Takeuchi and Higashiyama 2016).

The discovery of different types of attractants has further refined this concept. The AtLURE1-PRK6 signaling pathway promotes guidance of conspecific pollen tubes and accelerates the emergence of *A. thaliana* pollen tubes from the septum, providing a competitive advantage over heterospecific pollen tubes, such as those from *A. lyrata* (Zhong et al. 2019). In contrast, the evolutionarily conserved XIUQIU peptides attract pollen tubes from a wider range of Brassicaceae species and function independently of PRK6 (Zhong et al. 2019). Thus, ovule guidance utilizes both species-preferential and broadly acting signals, assuring efficient targeting of conspecific pollen while retaining some attraction to heterospecific pollen. Consequently, pollen tube guidance represents another molecular checkpoint that can restrict reproductive compatibility.

Even if heterospecific pollen tubes reach the ovule, they may face another prezygotic barrier at the synergid cells during PTR, the process culminating in pollen tube rupture and sperm release for double fertilization. Successful PTR involves a coordinated sequence: after the pollen tube contacts the receptive synergid cell, its growth slows, the synergid undergoes degeneration, and the pollen tube ultimately ruptures to release its sperm cells (Sandaklie-Nikolova L, et al. 2007; Leydon et al. 2015; Zhong et al. 2025). Before this transition, an autocrine signaling system maintains pollen tube integrity. Pollen tube-derived RALF4 and RALF19 signal through the ANXUR1/2 (ANX1/2) and BUDDHA’S PAPER SEAL1/2 (BUPS1/2) receptor complexes, along with the GPI-anchored co-receptors LLG2/3, to prevent premature rupture during pollen tube growth (Ge et al. 2017; Ge et al. 2019). As the pollen tube approaches the female gametophyte, ovule-derived RALF34 is proposed to compete with RALF4/19 for the ANX–BUPS receptor system, shifting the pollen tube from an integrity-maintaining state toward rupture (Ge et al. 2017; Feng et al. 2019). Thus, similar to the stigma, a signaling state maintained by an autocrine ligand can be redirected by a signal from the reproductive partner.

In parallel, the arriving pollen tube activates a female signaling pathway in the synergid cell. The CrRLK1L receptors FER, HERCULES RECEPTOR KINASE 1 (HERK1), and ANJEA (ANJ), along with the GPI-anchored proteins LORELEI (LRE) and ENODLs (ENs), function at the filiform apparatus and are required for compatible PTR (Escobar-Restrepo et al. 2007; Capron et al. 2008; Tsukamoto et al. 2010; Hou et al. 2016; Galindo-Trigo et al. 2020). Pollen tube-derived RALFs signal to this female machinery. RALF6, RALF7, RALF16, RALF36, and RALF37, regulated by the pollen tube-expressed transcription factors MYB97, MYB101, and MYB120, are essential for normal PTR; their combined loss leads to pollen tube overgrowth. These peptides interact with FER, ANJ, and HERK1, promoting receptor-complex assembly (Zhong et al. 2022). RALF4 and RALF19 can also function as pollen tube-derived ligands for FER–LRE, linking pollen tube arrival to downstream Ca²⁺ signaling (Gao et al. 2022). Pollen tube arrival induces FER-dependent relocalization of NORTIA (NTA) to the filiform apparatus, and FER, LRE, and NTA form a receptor–channel module through which pollen tube-derived RALFs stimulate Ca²⁺ responses in the synergid cell (Ju et al. 2021; Gao et al. 2022). Additionally, female-derived signals can act directly on the pollen tube to promote its rupture. In maize, the synergid-secreted defensin-like peptide ZmES4 activates the pollen tube K+ channel KZM1, causing membrane depolarization and ionic changes that rapidly lead to pollen tube rupture (Amien et al. 2010). Collectively, these signaling events couple perception of the arriving pollen tube to synergid responses and ultimately to pollen tube rupture and sperm release.

Despite this coordinated signaling process, heterospecific pollination can still fail. In interspecific crosses within *Rhododendron*, pollen tube overgrowth has been observed (Williams et al. 1982), and a similar phenotype occurs when *A. thaliana* ovules are pollinated with *A. lyrata* or *Cardamine flexuosa* pollen (Escobar-Restrepo et al. 2007; Müller et al. 2016). Several female-side factors have been implicated in this barrier. FER was initially associated with heterospecific PTR when *A. lyrata* and *C. flexuosa* pollen tubes exhibited a *fer*-like overgrowth phenotype in wild-type *A. thaliana* ovules (Escobar-Restrepo et al. 2007). Subsequent genetic evidence demonstrated that loss of FER further compromises reception of *A. lyrata* pollen tubes, supporting a direct role for FER-dependent signaling in heterospecific PTR (Ju et al. 2021). Notably, heterospecific pollen tubes can trigger FER/LRE-dependent Ca²⁺ oscillations in synergid cells yet still fail to complete PTR, indicating that activation of this signaling response alone is insufficient for successful reception (Ponvert and Johnson 2024). Species-dependent compatibility can also act at the final rupture step: ZmES4 rapidly induces rupture in maize pollen tubes, acts more slowly on pollen tubes of the related grass *Tripsacum dactyloides*, and fails to trigger rupture in more distantly related species, suggesting that compatibility between female-derived rupture signals and their pollen tube targets contributes to prezygotic reproductive isolation (Amien et al. 2010). ARTUMES (ARU), an OST3/6 subunit of the oligosaccharyltransferase complex involved in protein N-glycosylation, is particularly required for efficient reception of *A. lyrata* pollen tubes by *A. thaliana* ovules, with only a minor effect on conspecific PTR (Müller et al. 2016). Thus, core components of the conspecific reception machinery, female factors with more pronounced roles in heterospecific interactions, and species-dependent compatibility between female-derived rupture signals and their pollen tube targets can influence heterospecific PTR.

In spite of these advances, it remained unclear whether evolutionary variation in a key female PTR component could influence its compatibility with pollen tubes from other species. LRE provides an ideal system to investigate this question, as it is essential for conspecific PTR and belongs to a family of signaling co-receptors that has undergone evolutionary diversification. Our previous studies established LRE’s role in synergid-mediated PTR and provided the genetic and molecular tools necessary to examine its function (Tsukamoto et al. 2010; Liu et al. 2016; Noble et al. 2022).

Gene duplication, such as that resulting from whole-genome duplication, can allow duplicated genes to evolve differences in regulation or function because selective constraints change after duplication (Ramsey and Schemske 1998; Panchy et al. 2016). Extensive evolutionary analysis of the *LLG* family further supports focusing on LRE (Noble et al. 2022). The *A. thaliana* genome contains four *LLG* family members, each having distinct expression patterns: *LRE* is primarily expressed in the female gametophyte, *LLG2* and *LLG3* in the male gametophyte, and *LLG1* mainly in sporophytic tissues (Tsukamoto et al. 2010; Feng et al. 2019; Ge et al. 2019). Phylogenetic studies indicate that *LRE* and *LLG1* are closely related paralogs within Brassicaceae, and functional complementation experiments have shown a high degree of interchangeability among LLG-family proteins when expressed in the same cellular context. Thus, these genes are retained primarily because of expression differences rather than distinct protein functions (Noble et al. 2022).

Our previous phylogenetic analysis also revealed an interesting feature: the branch leading to *LRE* exhibited greater sequence divergence than that leading to *LLG1* (Noble et al. 2022). This raised the possibility that, despite maintaining its conserved role in PTR, *LRE* may have acquired sequence changes with additional functional effects. However, it remained unclear whether *LRE* and *LLG1* experienced different selective constraints after duplication, which sequence changes were associated with this divergence, and whether variation within *LRE* influences its function during interactions with heterospecific pollen tubes.

In this study, we examine these questions by investigating LRE function during heterospecific PTR within an evolutionary framework. We assess whether LRE and its co-receptor FER contribute to heterospecific PTR, whether LRE orthologs from different Brassicaceae species differ in their ability to support heterospecific PTR, and whether these functional differences are linked to evolutionary diversification within the LRE lineage. Additionally, we examine the effects of variation at evolutionarily significant LRE residues and determine whether variants with different PTR effects share predicted conformational characteristics. Collectively, these approaches permit us to evaluate whether diversification of a conserved female PTR component underlies species-dependent reproductive compatibility.

## Results

### Loss of LRE and FER negatively impacts heterospecific pollen tube reception in a pollen-species-dependent manner

The PTR constitutes a prezygotic barrier in interspecific crosses between *A. thaliana* and either *A. lyrata* or *Cardamine flexuosa* (Escobar-Restrepo et al. 2007; Müller et al. 2016), both of which belong to Lineage I of the Brassicaceae (Beilstein et al. 2006). Whether a similar PTR barrier occurs when *A. thaliana* is pollinated with the more distantly related Lineage II Brassicaceae species *Sisymbrium irio* has not been examined. To compare reproductive outcomes among these crosses, we pollinated *A. thaliana* pistils with *A. thaliana*, *A. lyrata*, or *S. irio* pollen and assessed pollen tube behavior using aniline blue staining (Fig. 1A-F).

**Figure 1.**
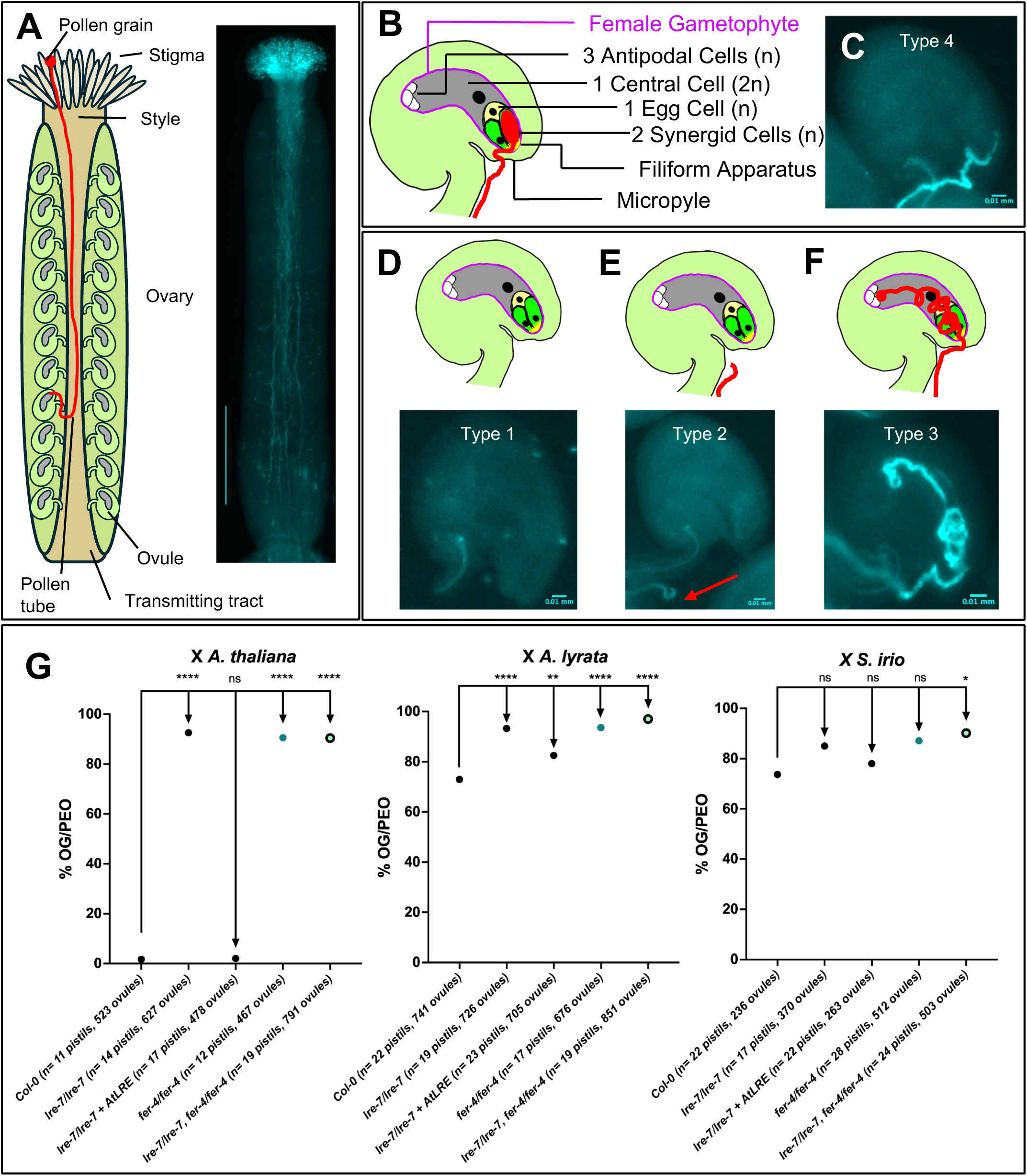
Visualization of progamic phase and associated defects, and quantification of pollen tube overgrowth mutant crosses to interspecific pollen. (A) Pollen tube growth in an *A. thaliana* pistil visualized by staining it with aniline blue. (Left) Diagram of *A. thaliana* pistil with pollen tube shown in red and different parts of the pistil are marked. (Right) An epifluorescent image of an aniline blue-stained pollinated pistil. Scale bar= 0.5 mm. (B and C) Diagram (B) and an aniline-blue stained epifluorescent image of pollen tube reception by an *A. thaliana* ovule (C, type 4). In the diagram (B), the pollen tube is marked in red and the persistent synergid cell is marked in green. The receptive synergid cell is filled with red, pointing to the successful discharge of the pollen tube cytoplasm into it and marking the completion of pollen tube reception by that ovule. In C, successful pollen tube reception and burst is inferred when the callose-rich pollen tube enters the ovule at the micropylar end and ceases growth in the area of the ovule proximal to the micropyle just after it enters the ovule. Neither the pollen tube cytoplasm or sperm cells are stained by DABS and so the bolus left by the burst pollen tube in the receptive synergid cell is not observed in this assay. (D-F) Defects in pollen tube-ovule interactions. Abnormal pollen tube growth can be divided into three other categories: the pollen tube fails to exit the transmitting tract and target an ovule (D, type 1), the pollen tube targets an ovule but does not enter the ovule (E, type 2), and the pollen tube targets the ovule and enters but does not stop to burst at the synergid cells; instead, it continues to grow and coils inside the female gametophyte (F, type 3). (C-F) Scale bar= 0.01 mm. (G) Quantification of the average pollen tube overgrowth per pollen tube-entered ovule (%OG/PEO) in crosses between female parents listed on the X-axis and *A. thaliana, A. lyrata, or S. irio* pollen. The average % OG/PEO for each female was compared to the % OG/PEO of wild type Columbia (Col-0) ecotype. % OG/PEO indicates the percentage of ovules in a silique that experienced pollen tube overgrowth in the female gametophyte out of all the ovules that received a pollen tube. The number of pistils crossed, and ovules scored is listed for each female parent. Statistical significance was determined by first using an arcsine transformation of percentage data, followed by pairwise *t-*tests between Col-0 and tested mutants; ns> 0.05, *p<0.05, **p<0.01, ***p<0.001, and ****p<0.0001. In each experimental series, Col-0, *lre-7*, and *lre-7* + *AtLRE_pro_:AtLRE-cYFP* were crossed and scored in parallel with the indicated test lines; thus, the control values shown in this panel are the contemporaneous controls from the same experimental block and may differ from control values shown in other figures.

We classified ovules into four categories based on pollen tube behavior. Types 1 and 2 represented defects occurring before pollen tube entry into the female gametophyte (Fig. 1D and 1E). Pollen tubes that entered the micropyle but continued to grow and coil within the female gametophyte were classified as Type 3, yielding clear morphological evidence of failed PTR (Fig. 1F). Pollen tubes that entered the female gametophyte and terminated near the expected position of the synergid cells without further growth or coiling were classified as Type 4, consistent with a reception-like outcome (Fig. 1B and 1C; see Methods).

The distribution of these four pollen tube outcomes differed markedly among the three crosses (Figure S1). In the conspecific *A. thaliana × A. thaliana* cross, 75% of ovules exhibited a Type 4 reception-like outcome, whereas only 1% exhibited Type 3 PTR failure. Consistent with previous reports (Escobar-Restrepo et al. 2007; Müller et al. 2016), the *A. thaliana × A. lyrata* cross showed a pronounced PTR barrier: 66% of all ovules exhibited PTR failure, whereas only 21% exhibited a Type 4 outcome. In the previously uncharacterized *A. thaliana × S. irio* cross, *S. irio* pollen encountered considerable barriers before entry into the female gametophyte, with 61% and 14% of ovules classified as Types 1 and 2, respectively. Of all ovules, 17% exhibited PTR failure, and 8% exhibited a Type 4 outcome. Thus, both *A. lyrata* and *S. irio* pollen encounter reproductive barriers in *A. thaliana*, although the relative frequencies of defects occurring before ovule entry and during PTR differ substantially between the two heterospecific crosses.

Having established PTR failure in both heterospecific crosses, we next asked whether LRE and FER, core components of conspecific PTR, influence defect severity. Following Müller et al. (2016), we quantified PTR failure as the percentage of pollen tube-entered ovules (PEO) exhibiting overgrowth (OG) [%OG/PEO = Type 3/(Type 3 + Type 4) × 100]. This metric specifically reports morphologically evident PTR failure among ovules that received a pollen tube and does not include defects occurring before entry. Because Type 3 classification required clear continued growth and coiling after entry, %OG/PEO provides a stringent and conservative estimate of PTR failure among pollen tube-entered ovules (see Methods).

As a conspecific control, we used *A. thaliana* pollen to pollinate wild-type (Col-0), *lre-7* single mutant, *fer-4* single mutant, *lre-7 fer-4* double mutant, and *lre-7* single mutant pistils complemented with *AtLRE_pro_:AtLRE-cYFP* (Liu et al. 2016; Figure S2). As expected, wild-type (1.7%) and *AtLRE_pro_:AtLRE-cYFP* complemented line showed efficient PTR (2.1%), whereas the single mutants and the *lre-7 fer-4* double mutant exhibited elevated PTR failure (92.6%, 90.6%, and 90.4%, respectively; Fig. 1G), consistent with the established roles of LRE and FER in conspecific PTR. For these experiments, statistical comparisons were made to Col-0 wild type because the objective was to determine how loss of endogenous LRE and FER function affected PTR.

When *A. thaliana* pistils were pollinated with heterospecific *A. lyrata* pollen, approximately 73% of pollen tube-entered ovules exhibited PTR failure, consistent with previous reports (Escobar-Restrepo et al. 2007; Müller et al. 2016; Fig. 1G). Loss of LRE further increased this defect, with *lre-7* pistils exhibiting 93.3% pollen tube overgrowth, significantly higher than in wild-type pistils. *fer-4* and *lre-7 fer-4* pistils also showed significantly elevated pollen tube overgrowth with *A. lyrata* pollen, at 93.5% and 97%, respectively. The AtLRE-complemented *lre-7* line showed a lower frequency of pollen tube overgrowth than *lre-7* (82.5% OG/PEO), reversing the effects of the *lre-7* mutation, although the complemented line remained significantly different from wild type (Fig. 1G). PTR failure was also frequent when *A. thaliana* pistils were pollinated with *S. irio* pollen. In this cross, pollen tube overgrowth was modestly higher in *lre-7* (85%) and *fer-4* (87.1%) pistils than in wild type (73.7%), but these differences were not statistically significant. The *lre-7 fer-4* double mutant, however, exhibited significantly greater pollen tube overgrowth than wild type, with 90.2% OG/PEO (Fig. 1G). The *lre-7* complemented with AtLRE-cYFP showed overgrowth levels similar to wild type (78%); however, because *lre-7* itself was not significantly different from wild type in this cross, we did not interpret this as evidence of significant complementation. Together, these results show that LRE and FER limit heterospecific PTR failure, although the magnitude of their contribution depends on the pollen species.

### LRE ortholog identity influences heterospecific pollen tube reception

We next asked whether replacing AtLRE in *A. thaliana* with an LRE ortholog from the pollen-donor species could improve heterospecific PTR. We generated *AtLRE_pro_:AlyLRE-cYFP* and *AtLRE_pro_:SiLRE-cYFP* constructs (Figure S2) and introduced them into the *A. thaliana lre-7* background. For each construct, three independent homozygous lines containing a single transgene copy were analyzed, as determined by segregation analysis (Methods), and confirmed by ddPCR (Figure S3). In all transgenic lines, cYFP fluorescence localized to the filiform apparatus of synergid cells, like full-length AtLRE-cYFP (Figure S4; Liu et al. 2016). Thus, both orthologs localized to the expected site of LRE function during PTR.

Before testing these lines in heterospecific crosses, we determined whether AlyLRE and SiLRE retained the LRE activity required for conspecific PTR. When pollinated with *A. thaliana* pollen, all AlyLRE and SiLRE lines complemented the *lre-7* PTR defect and supported PTR at levels comparable to AtLRE (Fig. 2A; Liu et al. 2016). For analyses of these transgenic lines, we compared them to *lre-7* complemented with *AtLRE_pro_:AtLRE-cYFP* rather than to Col-0, which was used as the reference in Figure 1. This transgenic line provides the appropriate matched control for the *lre-7* genetic background, AtLRE promoter, and cYFP-tagged protein used in the test constructs. These results established that both transgenes are functional and that both orthologs retained substantial basal LRE activity, allowing us to assess whether they differed specifically in their ability to support heterospecific PTR.

**Figure 2:**
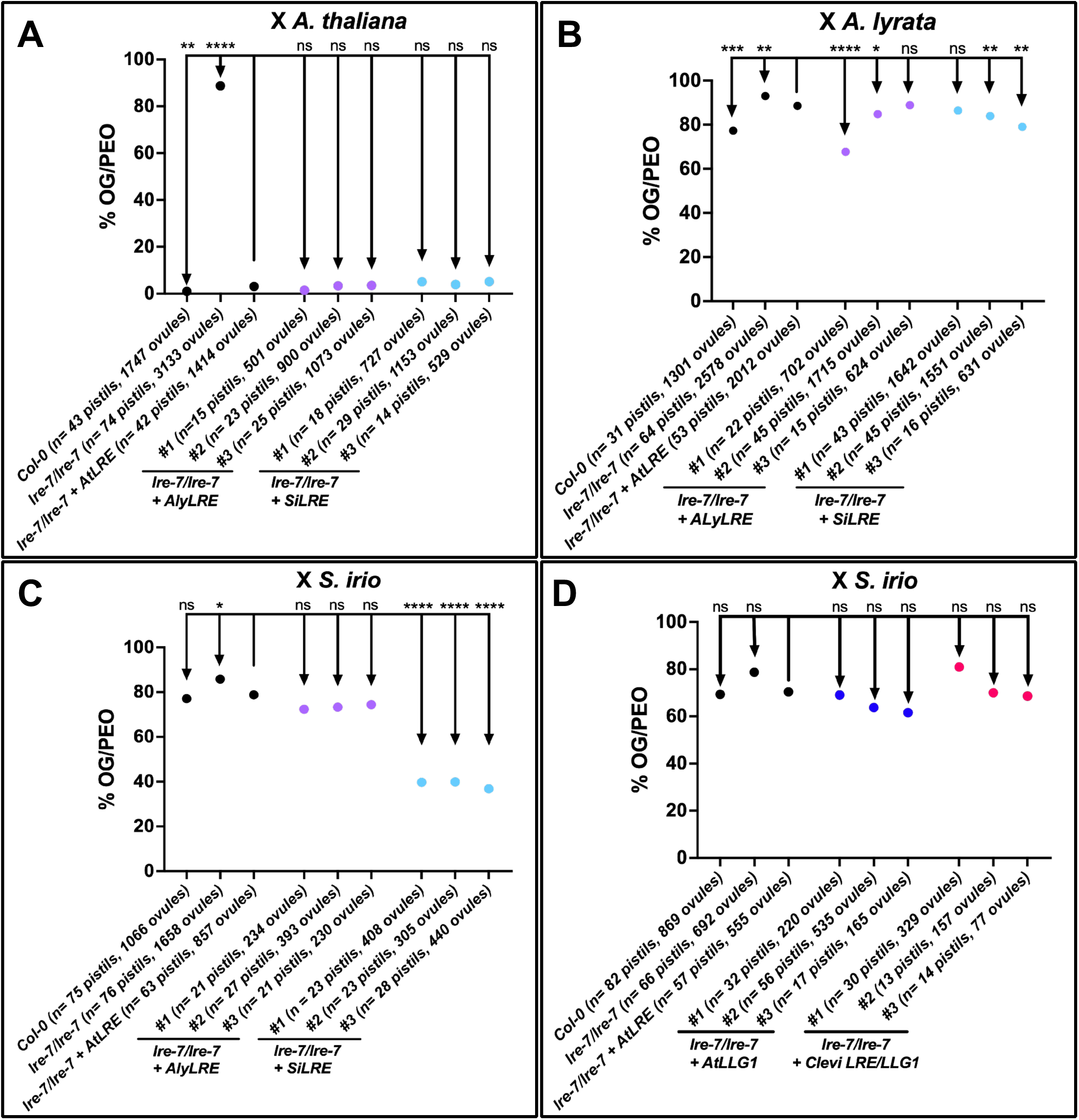
Full-length SiLRE significantly improves pollen tube reception in inter-lineage crosses to *S. irio* pollen. Quantification of the average pollen tube overgrowth per pollen tube-entered ovule (%OG/PEO) in crosses between female parents listed on the X-axis and *A. thaliana* pollen (A), *A. lyrata* pollen (B), and *S. irio* pollen (C, D). The average % OG/PEO for each female was compared to the % OG/PEO of *lre-7/lre-7 + AtLRE_pro_:AtLRE-cYFP*. In each construct, three independently transformed lines were crossed to *A. thaliana* pollen. % OG/PEO indicates the percentage of ovules in a silique that experienced pollen tube overgrowth in the female gametophyte out of all the ovules that received a pollen tube. The number of pistils crossed, and ovules scored is listed for each female parent. Statistical significance was determined by first using an arcsine transformation of percentage data, followed by pairwise *t-*tests between *lre-7/lre-7+ AtLRE_pro_:AtLRE-cYFP* and tested lines; ns> 0.05, *p<0.05, **p<0.01, ***p<0.001, and ****p<0.0001. In each experimental series, Col-0, *lre-7*, and *lre-7* + *AtLRE_pro_:AtLRE-cYFP* were crossed and scored in parallel with the indicated test lines; thus, the control values shown in this panel are the contemporaneous controls from the same experimental block and may differ from control values shown in other figures.

We first tested whether AlyLRE improved reception of *A. lyrata* pollen. Ovules expressing AtLRE and pollinated with *A. lyrata* pollen exhibited approximately 88.6% OG/PEO, whereas those expressing AlyLRE showed a modest but significant decrease to approximately 67.7% and 84.8% OG/PEO in two out of three lines tested (Fig. 2B). Ovules expressing SiLRE exhibited approximately 86.5%, 83.9%, and 79.1% OG/PEO in three lines when crossed to *A. lyrata* pollen. Although the first SiLRE line was comparable to AtLRE, the other two showed significantly lower OG/PEO when pollinated with A*. lyrata* pollen. Thus, some lines expressing either AlyLRE or SiLRE showed reduced OG/PEO, indicating that improved PTR was not restricted to the species-matched AlyLRE ortholog. In contrast, we observed a substantially larger effect with *S. irio* pollen. Ovules expressing AlyLRE did not significantly improve PTR relative to ovules expressing AtLRE and pollinated with *S. irio* pollen, with pollen tube overgrowth frequencies from 72.3% to 74.4% OG/PEO (Fig. 2C). By contrast, all three independent SiLRE lines showed markedly reduced pollen tube overgrowth (39.7%, 39.9%, and 36.8%) when pollinated with *S. irio* pollen (Fig. 2C). The improvement conferred by SiLRE also exceeded the level of PTR observed in wild-type *A. thaliana* pistils pollinated with *S. irio* (77.1%) (Fig. 2C). Independent experiments performed at the Institute of Experimental Botany, Czech Academy of Sciences, reproduced the SiLRE effect (Figure S5). Thus, expression of the species-matched SiLRE ortholog markedly improved reception of *S. irio* pollen among pollen tube-entered *A. thaliana* ovules.

We next asked whether this enhancement was a general property of *LRE/LLG* family members (Tsukamoto et al. 2010; Noble et al. 2022). We used previously described *A. thaliana lre-7* lines expressing full-length *A. thaliana LLG1* (*AtLRE_pro_:AtLLG1-cYFP*) or the single-copy *Cleome violacea LRE/LLG1* ortholog (*AtLRE_pro_:Clevi-LRE/LLG1-cYFP*) (Noble et al. 2022). Both fusion proteins localize primarily to the filiform apparatus when expressed in synergid cells and retain substantial activity during conspecific PTR (Noble et al. 2022). When these lines were pollinated with *S. irio*, none of the three independent lines expressing either construct significantly reduced pollen tube overgrowth relative to the AtLRE-complemented control (61.6%– 80.9% OG/PEO; Fig. 2D). Thus, the ability of SiLRE to improve reception of *S. irio* pollen is not a general property of the *LRE/LLG* family or of the ancestral single-copy LRE/LLG1-like protein, but instead depends on features that arose within the LRE lineage.

### The *LRE* clade, but not the *LLG1* clade, experienced relaxed selective pressures following duplication

AlyLRE and SiLRE affect heterospecific PTR modestly and variably with *A. lyrata* pollen, but strongly and consistently with species-matched SiLRE and *S. irio* pollen, suggesting that sequence changes acquired during LRE diversification may contribute to species-dependent heterospecific PTR. To identify candidate amino acid residues that might contribute to the species-matched enhancement of heterospecific PTR, we placed these functional differences in an evolutionary framework by reconstructing an expanded *LRE/LLG1* phylogeny and testing whether the two clades experienced different selective pressures following duplication.

We identified *LRE/LLG1* homologs from two Malvales and 25 Brassicales species, with broader sampling of non-Brassicaceae Brassicales, early-diverging Brassicaceae, and Lineage II Brassicaceae than in our previous analysis (Noble et al. 2022) (Table S1 and Table S2). The resulting phylogeny recovered two strongly supported Brassicaceae clades corresponding to *LRE* and *LLG1* (bootstrap support = 100% and 99%, respectively) (Figure S6). In contrast, the single-copy *LRE/LLG1* orthologs from non-Brassicaceae Brassicales fell outside both clades (Fig. 3; Figure S6). This pattern is consistent with *LRE* and *LLG1* having arisen from the Brassicaceae alpha whole-genome duplication (Noble et al. 2022). Both paralogs were retained in all 17 Brassicaceae species included in our sampling.

**Figure 3:**
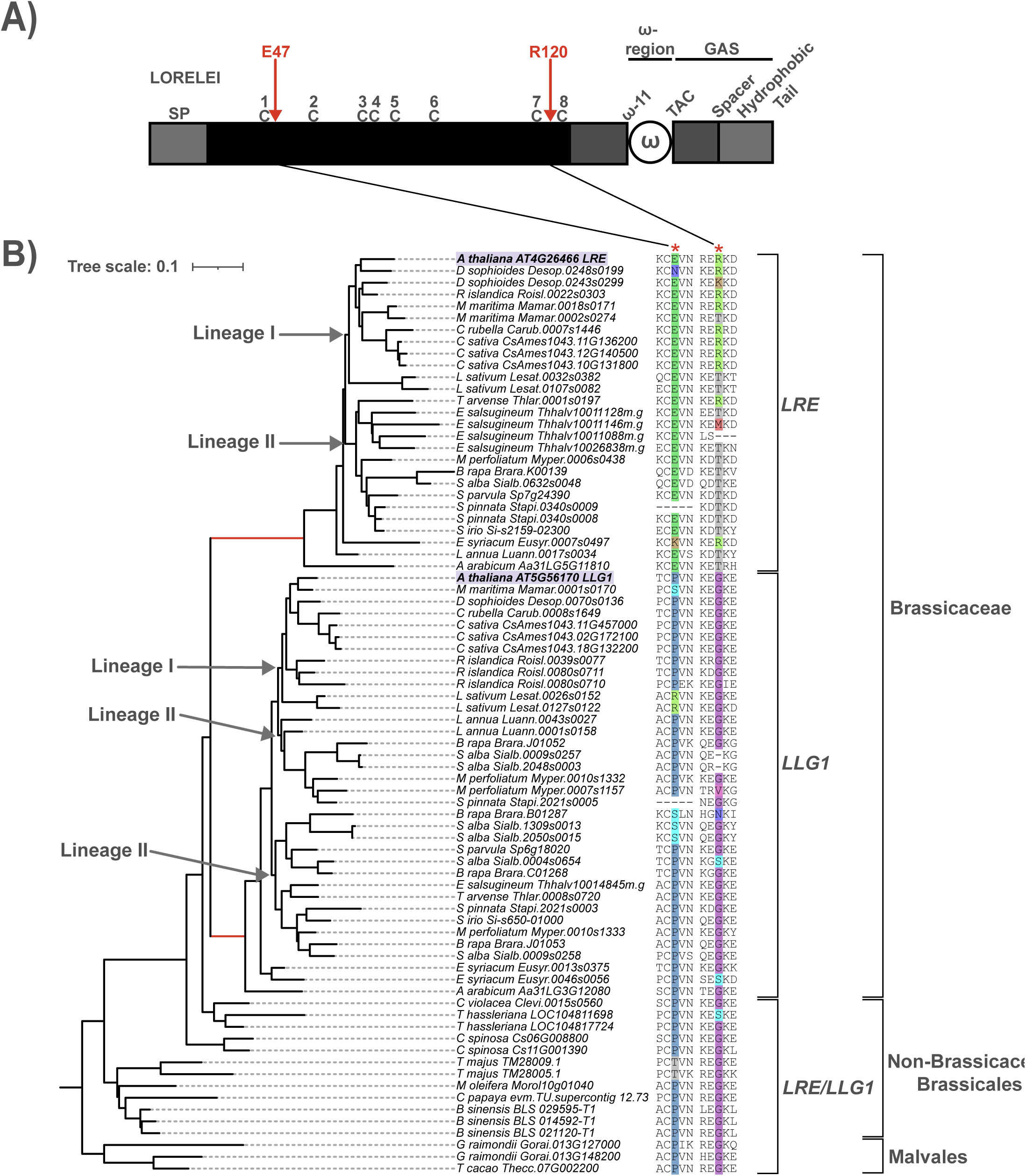
The *LRE* clade experienced a relaxation of purifying selection post-gene duplication in Brassicaceae. A) A diagram of the nascent *A. thaliana LRE* polypeptide with arrows indicating residues E47 and R120. The pre-processed protein contains a secretion peptide (SP), an extracellular domain (black) containing a modified eight-cysteine motif (M8CM; denoted by C1-8), a proline rich unstructured ω-11 region, ω, and a GPI-attachment signal (GAS) region. B) Maximum-likelihood phylogram inferred from 80 *LRE* and *LLG1* homologues retrieved from a set of 25 Brassicales and two Malvales species. The tree was rooted on Malvales *LRE/LLG1* homologues. *LRE* and *LLG1* clades from Lineage I and Lineage II Brassicaceae are marked with grey arrows. *LRE* and *LLG1* post-duplication branches are marked in red. *LRE* and *LLG1* from *A. thaliana* are highlighted with purple. *LRE* and *LLG1* duplicated early in the evolution of Brassicaceae. Sites potentially responsible for this signature of selection are shown at two short regions of the aligned polypeptide (corresponding to sites 45-49 and 118-122 of *A. thaliana LRE*); red asterisks denote E47 and R120.

The branch leading to the *LRE* clade was longer than the branch leading to *LLG1*, suggesting that the *LRE* clade experienced greater sequence divergence after duplication (Fig. 3). In the branch-model analysis, the *LLG1* clade did not differ significantly from the background lineages in selective pressure (p = 0.3853). In contrast, the *LRE* clade had a significantly higher nonsynonymous-to-synonymous substitution-rate ratio than the background (ω₁ = 0.37659 versus ω₀ = 0.28972; p = 0.0079; Table S3). Because both ω values remained below 1, the *LRE* clade was still under purifying selection, but that selection was significantly weaker than in the background lineages.

Site-model analyses also showed that selective constraint differed among sites within LRE/LLG1 proteins (*p* < 0.0001), with one class of sites under strong purifying selection (ω₀ = 0.19513; Table S3; Figure S7; Table S5). Likewise, Model C test identified a subset of sites that evolved faster in the *LRE* clade (ω₂,LRE = 0.53707) than in the background lineages (ω₂, background = 0.31928; *p* = 0.0077). However, the branch-site tests provided no clear evidence for positive selection on either the *LRE* post-duplication branch (*p* = 0.0838) or branches within the *LRE* clade (*p* = 1; Table S3). These results suggest that, after the duplication that gave rise to *LRE* and *LLG1*, selective constraint was reduced at a subset of sites in the *LRE* lineage, allowing greater sequence divergence while the protein remained under purifying selection. In contrast, *LLG1* showed no comparable shift in selective pressure and appears to have remained under constraints similar to those acting on the background lineages.

### Evolutionary analyses identify E47 and R120 as candidates for LRE functional diversification

Although the branch-site tests did not detect significant positive selection, the associated Empirical Bayes analyses identified two LRE sites of evolutionary interest. Glutamic acid at position 47 (E47) was assigned to an elevated-rate class along the LRE post-duplication branch and within the LRE clade, whereas arginine at position 120 (R120) was identified only within branches of the LRE clade (Table S4; Figure S7; Table S5). Because the branch-site tests were not significant, we considered these residues candidates for functional diversification rather than sites under positive selection. Notably, the residue homologous to LRE R120 in LLG1, glycine 120 (G120), contributes to RALF 23 binding specifically to LLG1 but not LRE (Xiao et al. 2019).

At the position corresponding to E47 in AtLRE, proline (P) predominated in the single-copy *LRE/LLG1* grade and the *LLG1* clade, indicating that P47 is the likely ancestral state (Fig. 3). By contrast, glutamic acid was present in nearly all sampled *LRE* orthologs, suggesting that a P-to-E substitution occurred before diversification of the *LRE* clade and was subsequently strongly conserved.

At the position corresponding to R120, glycine (G) predominated in the single-copy *LRE/LLG1* grade and the *LLG1* clade, indicating that G120 was likely ancestral. Within the LRE clade, however, glycine was largely replaced by arginine or threonine. Arginine predominated among Lineage I orthologs, whereas threonine predominated among Lineage II orthologs and occurred in selected additional lineages (Fig. 3).

E47 and R120 therefore represent two distinct patterns of LRE evolution: an ancestral substitution (E47) that became conserved throughout the LRE clade and a site (R120) that diversified in association with certain lineages. We therefore tested whether variation at these positions alters LRE’s ability to support heterospecific PTR.

### LRE variants retain basal activity, but R120T modestly compromises conspecific pollen tube reception

To test whether sequence changes associated with LRE divergence from *LLG1* contributed to LRE functional diversification, we replaced the *LRE* clade residues E47 and R120 with the corresponding ancestral, *LLG1*-like residues, P47 and G120. We generated the single-variant constructs *AtLRE_pro_:AtLRE(E47P)-cYFP* and *AtLRE_pro_:AtLRE(R120G)-cYFP* and the double-variant construct *AtLRE_pro_:AtLRE(E47P– R120G)-cYFP* (Figure S2). We also replaced the Lineage I-associated arginine at position 120 in AtLRE with threonine (T), the residue found predominantly at this position in Lineage II LRE orthologs, including in SiLRE, generating *AtLRE_pro_:AtLRE(R120T)-cYFP*. This variant allowed us to test whether lineage-associated variation at position 120 affects AtLRE in *A. thaliana’*s ability to support PTR with pollen from *S. irio*, a Lineage II species.

We introduced all four AtLRE variants into the *A. thaliana lre-7* background and selected three independent transgenic lines for each construct. cYFP fluorescence confirmed that all four variants were expressed in synergid cells and localized to the filiform apparatus, as in wild-type AtLRE-cYFP (Figure S4B to C, F to I). We first pollinated these lines with *A. thaliana* pollen to determine whether the substitutions affected the basal LRE activity required for conspecific PTR.

Lines expressing AtLRE-E47P or AtLRE-R120G exhibited low levels of pollen tube overgrowth (1.4%–10% OG/PEO) that were not significantly different from the AtLRE control, indicating that either ancestral-state substitution alone retained substantial conspecific PTR activity (Fig. 4A). The AtLRE-E47P–R120G double variant behaved similarly: all three independent lines exhibited very low levels of pollen tube overgrowth and were not significantly different from the AtLRE control (0.1%-0.7% OG/PEO; Fig. 4A). Thus, replacing E47 and R120 individually or together with their inferred ancestral, LLG1-like residues did not measurably compromise AtLRE function during conspecific PTR.

**Figure 4:**
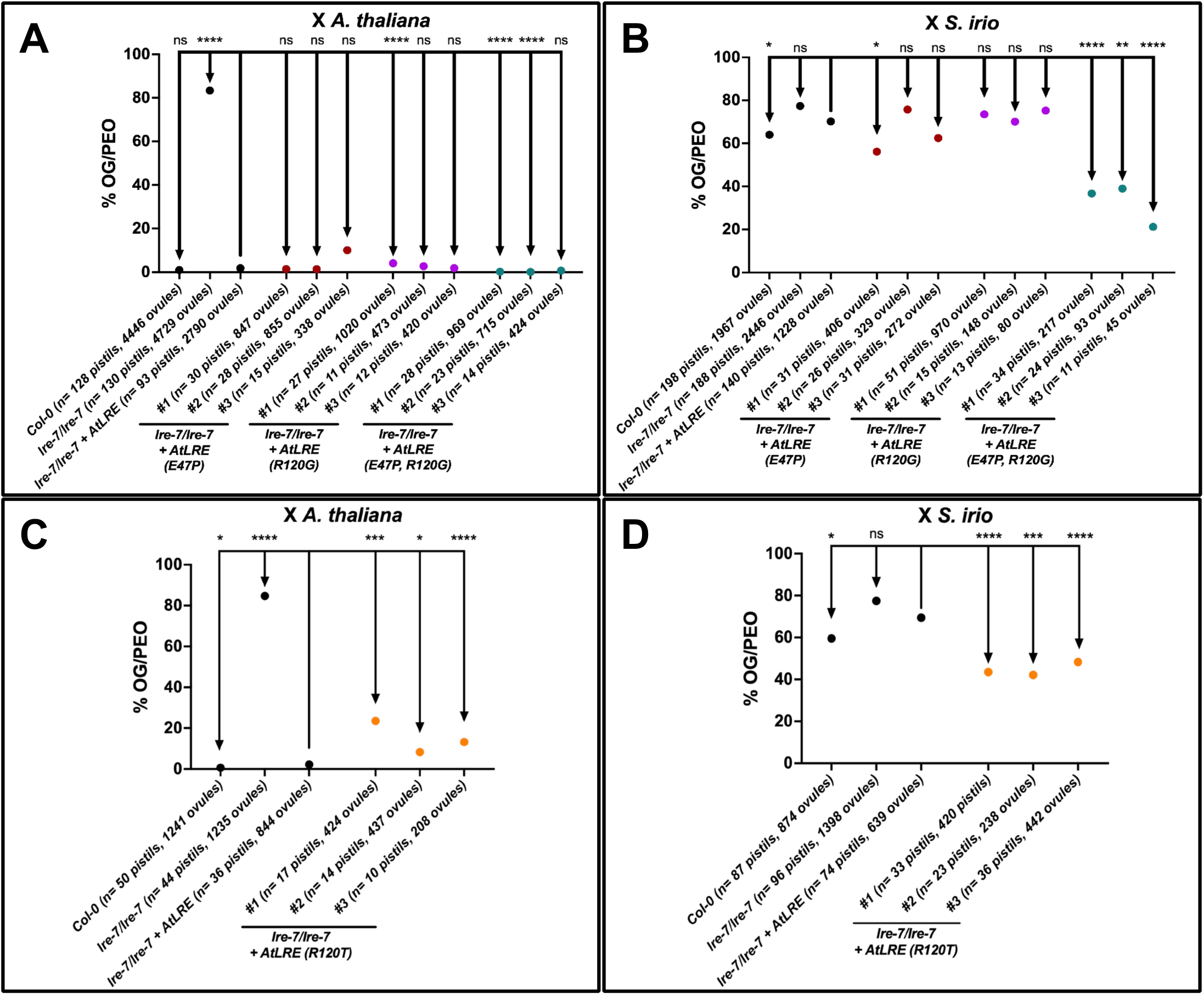
Both R120T single point mutant and E47P, R120G double point mutant improve pollen tube reception in inter-lineage cross to *S.irio* pollen. Quantification of the average pollen tube overgrowth per pollen tube-entered ovule (%OG/PEO) in crosses between female parents listed on the X-axis and *A. thaliana* pollen (A, C) and *S. irio* pollen (B, D). The average % OG/PEO for each female was compared to the % OG/PEO of *lre-7/lre-7 + AtLRE_pro_:AtLRE-cYFP*. In each construct, three independently transformed lines were crossed to *A. thaliana* pollen. % OG/PEO indicates the percentage of ovules in a silique that experienced pollen tube overgrowth in the female gametophyte out of all the ovules that received a pollen tube. The number of pistils crossed, and ovules scored is listed for each female parent. Statistical significance was determined by first using an arcsine transformation of percentage data, followed by pairwise *t-*tests between *lre-7/lre-7+ AtLRE_pro_:AtLRE-cYFP* and tested lines; ns> 0.05, *p<0.05, **p<0.01, ***p<0.001, and ****p<0.0001. In each experimental series, Col-0, *lre-7*, and *lre-7* + *AtLRE_pro_:AtLRE-cYFP* were crossed and scored in parallel with the indicated test lines; thus, the control values shown in this panel are the contemporaneous controls from the same experimental block and may differ from control values shown in other figures.

By contrast, substitution of R120 with the Lineage II-associated threonine significantly reduced conspecific PTR efficiency. All three AtLRE-R120T lines exhibited significantly higher pollen tube overgrowth than the AtLRE control, ranging from 8.3% to 23.5% OG/PEO (Fig. 4C). Nevertheless, these values remained substantially below the 84.6% overgrowth observed in *lre-7*, demonstrating that AtLRE-R120T retained considerable LRE activity. Thus, unlike the ancestral R120G substitution, introducing the Lineage II-associated R120T substitution partially compromised AtLRE’s ability to support reception of *A. thaliana* pollen tubes.

### Distinct LRE sequence changes improve reception of *S. irio* pollen

Because all four AtLRE variants retained substantial LRE activity during conspecific PTR, although R120T partially reduced reception of *A. thaliana* pollen, we next tested whether these substitutions altered heterospecific PTR in the *A. thaliana × S. irio* interlineage cross. In heterospecific crosses, SiLRE produced a substantially greater improvement with *S. irio* pollen than AlyLRE did with *A. lyrata* pollen (Fig. 2C). We therefore focused the current analysis of the LRE variants on the *A. thaliana × S. irio* cross, where the larger phenotypic effect provided greater resolution for determining whether candidate residues affect heterospecific PTR.

Reverting either residue individually to its inferred ancestral, LLG1-like state did not consistently improve reception of S*. irio* pollen. None of the three R120G lines differed significantly from *lre-7* complemented with wild-type AtLRE, and two of the three E47P lines likewise showed no improvement (Fig. 4B). Although one E47P line exhibited significantly less pollen tube overgrowth, the effect was modest and was not reproduced in the other independent lines. Thus, neither the E47P nor the R120G substitution alone was sufficient to enhance the ability of AtLRE expressed in *A. thaliana* ovules to support the reception of *S. irio* pollen.

By contrast, replacing the Lineage I-associated arginine at position 120 with the threonine present in SiLRE and predominant among Lineage II LRE orthologs (R120T) significantly improved PTR with *S. irio* pollen. All three R120T lines exhibited significantly less pollen tube overgrowth than the wild-type AtLRE control (69.5% OG/PEO), with the strongest line exhibiting 42.1% OG/PEO (Fig. 4D). This effect is particularly notable because the same R120T substitution significantly reduced PTR efficiency with conspecific *A. thaliana* pollen. Thus, R120T did not simply increase overall LRE activity; rather, it altered LRE function in a pollen-species-dependent manner, reducing compatibility with *A. thaliana* pollen while improving compatibility with *S. irio* pollen. These results show that changing a single AtLRE residue to the predominant state in the pollen-donor lineage is sufficient to shift heterospecific PTR toward greater compatibility with *S. irio* pollen.

Unexpectedly, the combined ancestral-state substitutions E47P–R120G also markedly improved reception of *S. irio* pollen. All three independent lines expressing the double variant exhibited significantly less pollen tube overgrowth than the wild-type AtLRE control, with values of 36.7%, 39.0%, and 21.2% OG/PEO (Fig. 4B). This reproducible effect was not observed when either ancestral-state substitution was introduced individually, indicating that the effects of residues 47 and 120 depend on sequence context. Notably, unlike R120T, the E47P–R120G variant improved reception of *S. irio* pollen without measurably compromising conspecific PTR with *A. thaliana* pollen.

Together, these results show that distinct sequence changes in AtLRE can improve reception of *S. irio* pollen. The Lineage II-associated R120T substitution produces a pollen-species-dependent shift in PTR, whereas the E47P–R120G combination improves heterospecific reception without a detectable loss of conspecific PTR activity. Thus, residues 47 and 120 contribute to AtLRE’s ability to support heterospecific PTR and identify sequence context within LRE as an important determinant of incompatibility with *S. irio* pollen.

### Pollen tube reception-enhancing LRE variants share predicted conformational features

Because R120T and E47P–R120G both improved the reception of *S. irio* pollen despite containing distinct mutations, we asked whether these variants shared structural conformational features that distinguished them from variants that did not consistently improve heterospecific PTR. We first generated standard AlphaFold 3 predictions for wild-type AtLRE, E47P, R120G, R120T, and E47P–R120G using a randomized seed approach. These predictions did not display a consistent structural difference that clearly delineated the variants from wild-type AtLRE or from one another (Figure S8). Prediction confidence was lowest in the N- and C-terminal regions, prompting us to examine whether conformational sampling differed among variants in these regions.

We therefore used subsampled AlphaFold, which increases the diversity of predicted conformations by reducing the depth of the multiple-sequence alignment used for modeling (Monteiro da Silva et al. 2024). Across all variants, the central α-helical bundle remained well conserved, whereas the N- and C-terminal regions displayed greater conformational variability and lower prediction confidence, likely due to poor sequence coverage in the multiple-sequence alignment (Figure S9).

Comparison of the highest-ranked structures revealed that R120T and E47P– R120G adopted predicted conformations in which the N- and C-terminal regions were farther apart. By contrast, wild-type AtLRE and R120G adopted more compact structural ensembles in which the termini were closer together, while E47P exhibited an intermediate arrangement. The R120T and E47P–R120G (<u>EPRG in</u> Fig. 5 and S10) ensembles also sampled broader conformational spaces than those of wild-type AtLRE, R120G, or E47P (Fig. 5A).

**Figure 5:**
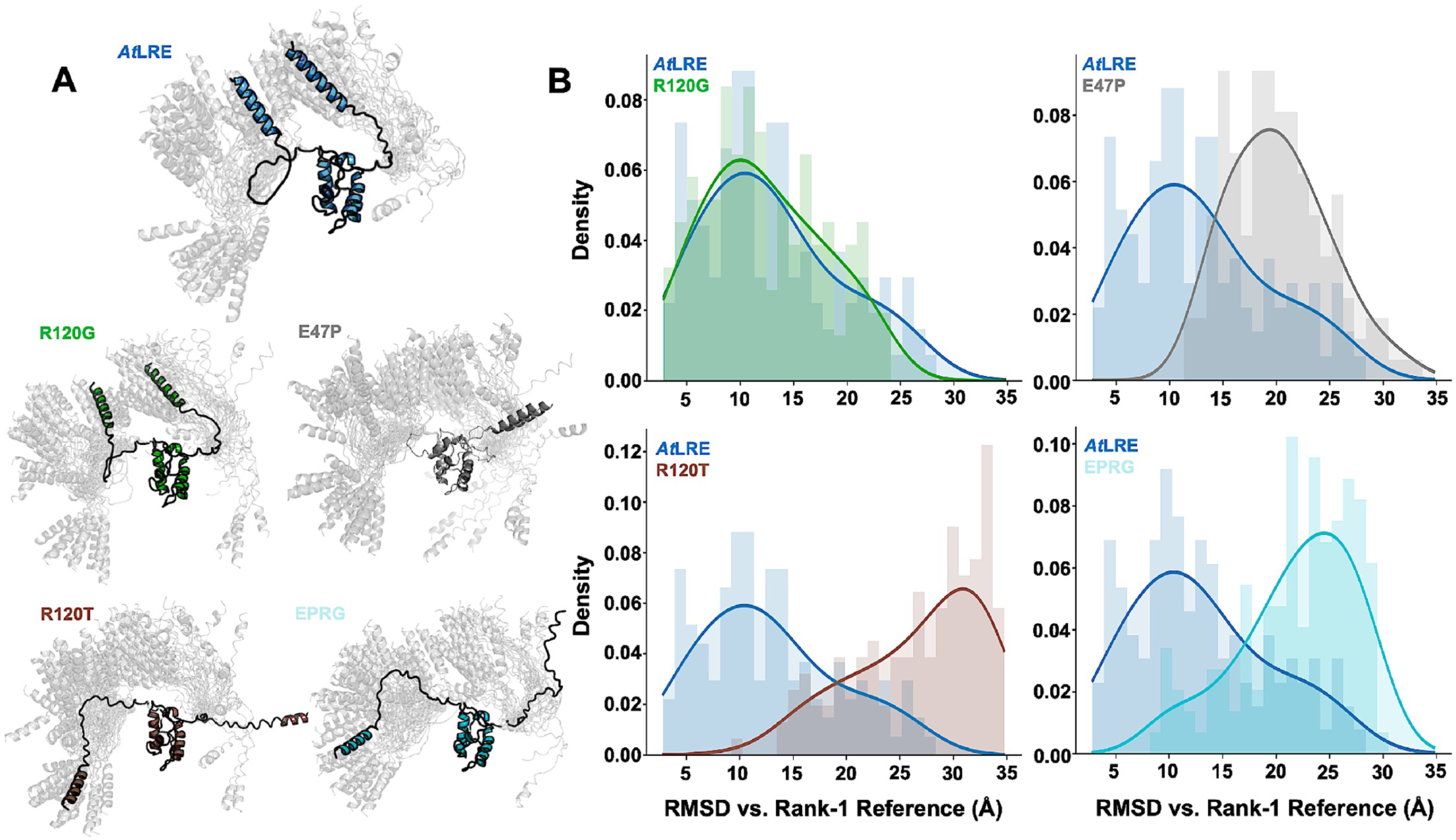
Structural models from subsampled AlphaFold suggest distinct N- and C-terminal ensembles between LRE variants. (**A**) Renderings of the fifty highest ranked structures (shown in grey) from each conformational ensemble. The highest ranked structures are highlighted in color for WT (blue), R120G (green), E47P (grey), R120T (brown) and E47P-R120G (EPRG, cyan). (**B**) RMSD plots of each variant, calculated using the highest ranked structural model as a reference. Variants are also compared to WT *At*LRE using same coloring as (**A**).

RMSD analysis within each ensemble showed similar results. R120T and E47P– R120G had the widest spread relative to their top-ranked structures, showing more variability across the predicted models. E47P showed less variation, and R120G was closest to wild-type AtLRE (Fig. 5; Figure S10).

The shared predicted features of R120T and E47P–R120G parallel their shared ability to improve the reception of *S. irio* pollen. Although these computational evaluations do not establish the underlying molecular mechanism, they suggest that distinct amino acid substitutions at positions 47 and 120 may relieve the heterospecific PTR barrier by altering the conformational states of LRE samples. Together, our genetic, evolutionary, and structure prediction results implicate these residues in limiting heterospecific *S. irio* PTR by *A. thaliana* ovules and provide a testable structural hypothesis for how LRE sequence variation can modulate prezygotic reproductive compatibility.

## Discussion

### LRE contributes to lineage-associated compatibility during heterospecific pollen tube reception

Our results show that LRE, a core component of conspecific PTR, also influences compatibility between heterospecific pollen tubes and the female reception machinery, thus affecting whether PTR succeeds or fails. This indicates that LRE can influence heterospecific compatibility. Several female components have previously been implicated in heterospecific PTR. FERONIA (FER), a core component of conspecific PTR, was initially linked to heterospecific reception when *A. lyrata* and *Cardamine flexuosa* pollen tubes were found to overgrow within wild-type *A. thaliana* female gametophytes, producing a phenotype resembling that of *fer* mutants (Escobar-Restrepo et al. 2007). This observation led to the proposal that divergence in FER-dependent pollen tube– synergid signaling could contribute to reproductive isolation. More direct genetic evidence was subsequently provided by Ju et al. (2021), who showed that reception of *A. lyrata* pollen was severely compromised in *fer-1* ovules and that the enhancement of heterospecific reception by filiform apparatus-targeted NTA remained dependent on FER. Thus, components of the core machinery that mediate conspecific PTR could also influence the outcome of heterospecific interactions.

ARU, an OST3/6 subunit involved in protein N-glycosylation, provides a different example. Loss of ARU strongly increases overgrowth of *A. lyrata* pollen tubes while leaving conspecific PTR largely unaffected (Müller et al. 2016). The greater sensitivity of heterospecific reception to ARU function suggests that proper configuration of the female reception machinery is especially important when interacting male and female components are less well matched. Thus, the heterospecific PTR barrier can arise through altered performance of molecular components whose functions are embedded within the normal female reception machinery rather than through a dedicated pathway for rejecting foreign pollen.

Our results expand this framework by showing that sequence variation within LRE modulates heterospecific compatibility in a manner that appears to reflect evolutionary lineage rather than strict species matching. AlyLRE did not uniquely improve reception of *A. lyrata* pollen; both AlyLRE and SiLRE produced modest and variable improvements in some lines. By contrast, with the more distantly related Lineage II species *S. irio*, SiLRE consistently and markedly improved reception, whereas AlyLRE did not improve reception relative to AtLRE. Together with the effects of substitutions at residues 47 and 120, this evidence indicates that lineage-associated sequence diversification within LRE can alter the compatibility of the female reception machinery with pollen tubes from different evolutionary lineages. The results therefore do not support a simple species-specific recognition system in which each LRE ortholog preferentially receives pollen from its own species. Rather, variation within this conserved PTR component appears to modify the compatibility of the female machinery with heterospecific pollen in a lineage-associated and sequence-context-dependent manner. Collectively, these results indicate that diversification of LRE following duplication generated protein-level variation that contributes to lineage-associated reproductive compatibility.

These conclusions should, however, be interpreted considering what our assay directly measures. PTR success was inferred from pollen tube morphology in aniline blue-stained pistils. This readout provides a strong and tractable means of distinguishing PTR failure, indicated by continued pollen tube growth and coiling after micropyle entry, from a reception-like outcome, indicated by pollen tube termination near the synergid cells region in ovules. However, it does not directly demonstrate downstream hallmarks of productive reception, including sperm discharge and initiation of double fertilization. This distinction may be especially relevant in heterospecific pollinations, in which pollen tube termination could, in principle, reflect growth arrest without sperm discharge. We attempted to establish *S. irio* lines carrying fluorescent pollen- and sperm-cell markers that would permit direct visualization of these events, but despite repeated efforts, we were unable to transform *S. irio* successfully. Consequently, although the reduced overgrowth observed with SiLRE and the AtLRE variants R120T and E47P–R120G demonstrates that LRE sequence variation shifts heterospecific pollen tube behavior toward a PTR-like outcome, whether these changes also increase sperm discharge and productive fertilization remains to be determined. Future development of transformation methods or alternative live-cell markers for *S. irio* will be important for directly linking these changes in pollen tube behavior to fertilization outcomes. Nevertheless, the consistent reductions in pollen tube overgrowth seen across independent transgenic lines provide strong comparative evidence that LRE sequence variation is associated with protein-level variation that may contribute to lineage-associated reproductive compatibility.

### Evolutionary diversification of LRE identifies residues 47 and 120 as candidates for functional divergence

Consistent with the idea that diversification of LRE contributed to its species-dependent behavior during heterospecific PTR, our expanded evolutionary analyses, building on our previous work (Noble et al. 2022), revealed greater sequence divergence in *LRE* than in its paralog *LLG1* following their duplication. Our earlier studies suggested that retention of the LLG family was driven largely by regulatory subfunctionalization: family members retained substantial functional interchangeability when expressed in the same cellular context but acquired distinct expression domains (Noble et al. 2022). The present results suggest an additional layer of divergence at the protein level. Although both *LRE* and *LLG1* have been broadly retained across Brassicaceae, purifying selection was significantly relaxed in the *LRE* clade, particularly at a subset of sites, whereas selective pressure on *LLG1* remained like that on the background lineages (Table S3). One possible explanation is that the relatively restricted expression of *LRE* in the female gametophyte reduced pleiotropic constraints on its sequence, permitting greater evolutionary flexibility, whereas the varied developmental functions of *LLG1* may have imposed stronger constraints on sequence divergence. Thus, our data extend the previous model of predominantly expression-based divergence within the LLG family by showing that LRE also underwent protein-level functional diversification that contributes to species-dependent compatibility during heterospecific PTR.

Within this pattern, evolutionary analyses highlighted E47 and R120 as residues of particular interest (Table S4; Table S5; Figure S7). Our results showed that E47 shows a sequence change associated with the early divergence of LRE from LLG1, whereas R120 captures subsequent lineage-associated diversification within the LRE clade (Fig. 3). The structural and functional contexts of these residues provided additional reasons to consider their contributions to LRE function. E47 lies immediately after the first cysteine in LRE and is predicted by AlphaFold to reside in a loop following an α-helix (Jumper et al. 2021; Fleming et al. 2025). R120 has an independently established functional context: the homologous residue in LLG1 contributes to RALF peptide-binding specificity, and substitutions at this position can alter RALF binding (Xiao et al. 2019). Thus, both residues occupy positions that could influence LRE function, although their separate evolutionary histories suggest they may have played different roles in its diversification.

The functional consequences of variation at these positions were also informative. Replacing E47 or R120 individually with their ancestral, LLG1-like residues did not substantially impair conspecific PTR, and the combined E47P–R120G variant likewise retained normal basal LRE activity. This result agrees with our previous observation that LLG1, which has proline and glycine at the equivalent positions, can rescue the *lre-7* PTR defect (Noble et al. 2022). In contrast, changing the Lineage I-associated R120 residue to the Lineage II-associated threonine caused only a modest decrease in conspecific reception of *A. thaliana* pollen, indicating that the R120T substitution retains much of LRE function. These data suggest that considerable sequence variation at positions 47 and 120 can be accommodated without loss of core LRE function, although the Lineage II-associated R120T substitution modestly reduces conspecific PTR efficiency.

### Effects of E47 and R120 substitutions on heterospecific pollen tube reception

Although substitutions at E47 and R120 retained substantial basal LRE function, their effects during heterospecific pollination suggest that these residues contribute to reproductive compatibility rather than to LRE function alone. The strongest evidence for a lineage-associated effect was obtained with the R120T substitution, which substantially enhanced the PTR of *S. irio* pollen. Nevertheless, position 120 cannot by itself explain species-dependent compatibility. Full-length SiLRE produced a greater improvement in reception of *S. irio* pollen than the R120T substitution alone, indicating that additional sequence features of SiLRE add to its species-matched activity. This interpretation is also supported by the conspecific pollination results. The R120T substitution modestly reduced reception of *A. thaliana* pollen, whereas full-length SiLRE supported conspecific PTR as well as AtLRE. This suggests that the effect of T120 depends on the sequence context in which it occurs, with other residues in SiLRE likely modifying or compensating for its effect.

The E47 results add an important dimension to this interpretation. E47P alone did not consistently improve reception of *S. irio* pollen, nor did the ancestral R120G substitution. Remarkably, however, combining these two individually ineffective substitutions in E47P–R120G substantially improved heterospecific PTR in all three independent lines while leaving conspecific reception essentially unaffected. The strong phenotype of the double variant therefore implicates E47 in a context-dependent interaction with residue 120. It demonstrates that improved heterospecific PTR cannot be explained solely by matching residue 120 to the pollen-donor Lineage. Instead, positions 47 and 120 together influence the incompatibility imposed by wild-type AtLRE during reception of *S. irio* pollen. Together, these data indicate that LRE-dependent reproductive compatibility is influenced by residue identity and sequence context. In contrast, LRE sequence variation represents only one component of a wider species-dependent signaling interaction.

### LRE may mediate heterospecific compatibility through interactions with pollen tube-derived signals

Although changes in LRE substantially weakened the heterospecific PTR barrier, they did not fully restore reception, suggesting that LRE-dependent compatibility operates within female signaling machinery and that matching additional female signaling components to the pollen donor could further improve PTR. FER is an obvious candidate because it functions with LRE during PTR, and our genetic results show that loss of FER further compromises reception of *A. lyrata* pollen. In *A. thaliana* synergid cells, the CrRLK1L receptors ANJEA and HERK1 also function redundantly with FER, and LRE, FER, ANJEA, and HERK1 can associate in a multimeric receptor complex (Galindo-Trigo et al. 2020). Additional female components that could be tested include the GPI-anchored EARLY NODULIN-LIKE proteins (ENODLs), which interact with FER and are also required for efficient PTR (Hou et al. 2016).

Compatibility is also likely to depend on the pollen tube-derived signals to which this machinery responds. In our experiments, the female genetic background remained *A. thaliana,* while pollen identity varied, and changing LRE sequence altered PTR in a pollen-species-dependent manner. We therefore propose that LRE sequence variation may influence how effectively the female LRE-associated receptor machinery responds to signals from the incoming pollen tube (Fig. 6). In this model, one or more *S. irio*-derived factors interact inefficiently with AtLRE, resulting in ineffective activation of the female reception machinery, whereas replacement of AtLRE with SiLRE improves compatibility with these signals and promotes productive PTR (Fig. 6).

**Figure 6:**
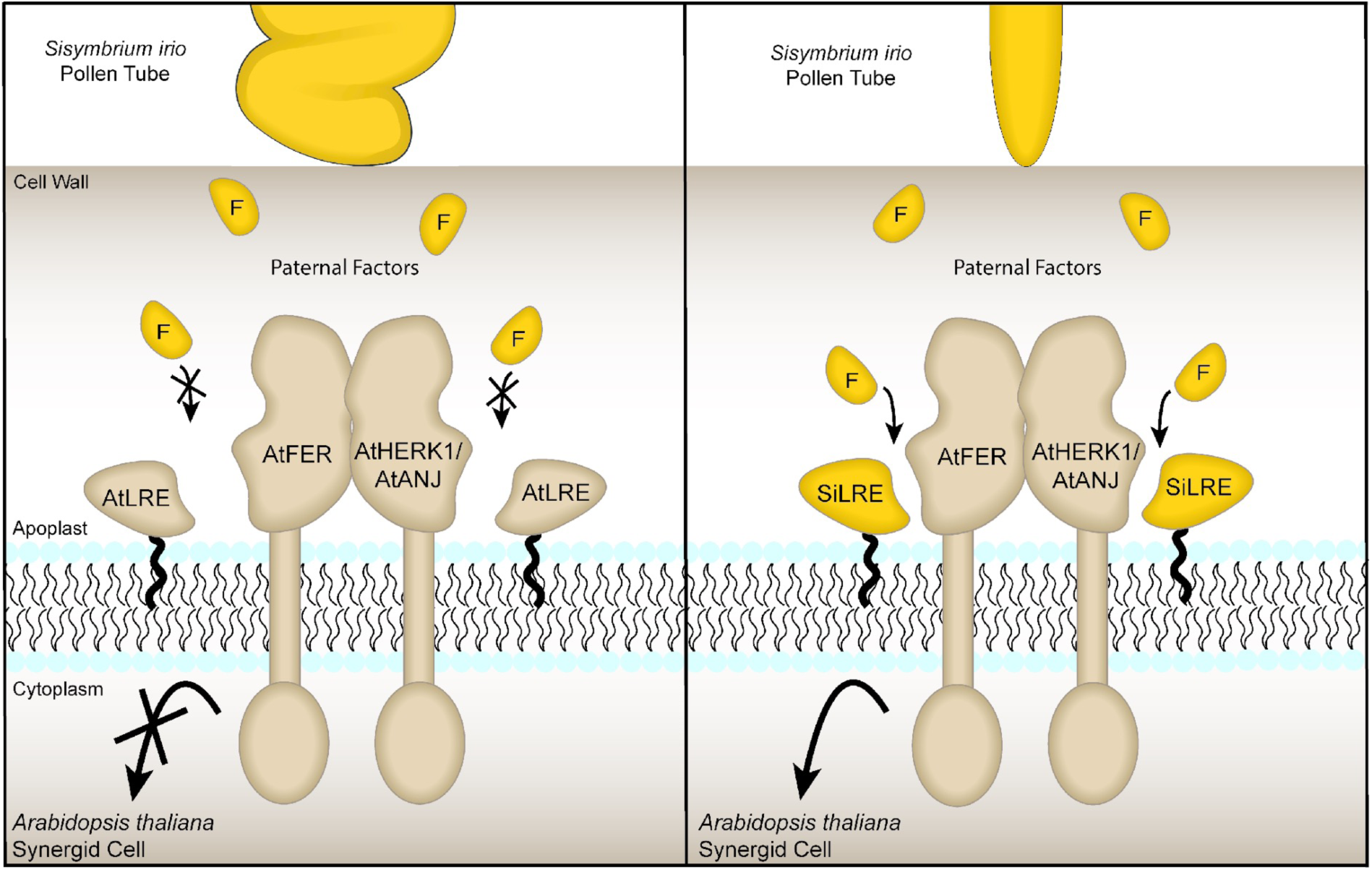
A working model of the interaction between *S. irio* pollen tubes and *A. thaliana* synergid cells in ovules with and without *S. irio* LRE. In a wild-type cross between *A. thaliana* and *S. irio*, we observe that pollen tube reception fails, possibly due to incompatible paternal factors (F) that the AtLRE-AtFER-AtHERK1/AtANJ complex does not recognize and therefore does not bind. When AtLRE is replaced with SiLRE in the synergid cells, we observe an increase in normal pollen tube reception with *S. irio* pollen tubes, potentially due to pollen tube recognition by SiLRE. In this scenario, the *S. irio* paternal factors (F) bind SiLRE and initiate SiLRE-AtFER-AtHERK1/AtANJ complex formation for downstream signaling leading to successful pollen tube reception.

RALF peptides are particularly strong candidates for these pollen tube-derived factors. A biochemical precedent comes from RALF23 signaling through LLG1: RALF23 binds LLG1, which contains glycine at the position homologous to LRE R120, whereas wild-type LRE, which contains arginine at this position, shows little or no detectable RALF23 binding; substitution of LRE R120 with the LLG1-like glycine restores measurable RALF23 binding (Xiao et al. 2019). Thus, the amino acid at this homologous position can contribute strongly to RALF-binding specificity within the LLG protein family, although the difference in affinity between LLG1 and LRE-R120G also indicates that additional sequence context is important. This relationship is especially relevant to our findings because variation at the same LRE position alters PTR in a pollen-species-dependent manner.

Independent genetic evidence further identifies biologically relevant pollen tube-derived RALFs that function during PTR. The pollen tube-expressed transcription factors MYB97, MYB101, and MYB120 are required for normal reception, and five of their downstream targets - RALF6, RALF7, RALF16, RALF36, and RALF37 - are required for efficient PTR. Their combined loss causes pollen tube overgrowth, and these peptides can interact with FER, ANJEA, and HERK1 and promote receptor-complex interactions (Zhong et al. 2022). These data make the five RALFs strong candidates for the pollen tube-derived factors represented schematically in Fig. 6. Together, this evidence supports a model in which LRE sequence diversification modulates the response of the female reception machinery to pollen tube-associated signals (Fig. 6). In this view, the prezygotic PTR barrier is not controlled by a single molecular gatekeeper but emerges from compatibility among interacting male and female signaling components.

### Reconstituting lineage-matched female receptor complexes to further enhance heterospecific pollen tube reception

A further prediction of this model is that LRE sequence variation is only one contributor to heterospecific compatibility, and that reception may improve further when additional interacting components of the female receptor machinery are also matched to the pollen-donor lineage. In our experiments, SiLRE and AtLRE variants weakened the *A. thaliana × S. irio* PTR barrier in an otherwise *A. thaliana* female background, implying that incompatibility can be partially relieved by tweaking a single component of the complex. However, because LRE functions within a multimeric receptor module that includes FER and the related CrRLK1L receptors ANJEA and HERK1 (Galindo-Trigo et al. 2020) and is probably shaped by additional associated factors such as ENODLs (Hou et al. 2016), it will be informative to ask whether “reconstituting” a more lineage-matched complex in *A. thaliana* improves compatibility beyond what is achieved by LRE alone. For example, expressing *S. irio* orthologs of FER and/or other core receptors together with SiLRE in *A. thaliana* synergid cells could test whether co-adapted receptor components provide additive or synergistic improvements in heterospecific PTR. If such a lineage-matched complex reconstitution yields a stronger reduction in pollen tube overgrowth and/or increases downstream indicators of successful PTR, it would support a model in which species-dependent PTR outcomes emerge from cumulative matching effects across male-female signaling components rather than from a single gatekeeping factor.

### Conclusions

Our results show that a component of the conserved machinery required for normal pollen tube reception can also shape reproductive compatibility between species. Rather than acting as a dedicated barrier to heterospecific pollen, LRE may influence how effectively the female reception machinery functions with pollen tubes from different evolutionary lineages. Importantly, this property can be altered by relatively limited changes in LRE sequence without eliminating its core role in conspecific PTR. The opposing effects of LRE orthologs and of substitutions at residues 47 and 120 additionally indicate that heterospecific compatibility is determined by sequence context rather than by a simple species-specific recognition code.

These results suggest that prezygotic barriers at the PTR stage may emerge through evolutionary divergence of otherwise conserved male-female signaling components. We propose that LRE sequence variation may modify the compatibility between pollen tube-derived signals and the female receptor machinery, with successful reception ultimately depending on both signal recognition and productive transmission through downstream PTR pathways. In this view, heterospecific PTR failure may reflect imperfect molecular matching across the signaling system rather than the action of a single barrier factor. Defining the pollen tube-derived signals involved and determining how their interactions with LRE-associated receptor complexes differ across lineages will be important to understand how reproductive barriers evolve and how they might eventually be modified to broaden interspecific compatibility.

## Methods

### Plant materials and growth conditions

*A. thaliana*, *Arabidopsis lyrata,* and *Sisymbrium irio* seeds were sterilized as follows: 0.05 - 0.1 mL of seeds were added to a 1.5 mL microcentrifuge tube and placed in a desiccator that contained an open beaker filled with 95 mL bleach and 5 mL HCl. After 3 hours, sterilized seeds were plated on MS plates with 2% sucrose and appropriate antibiotics for selection. *A. thaliana* seeds were stratified for 3 days in the dark at 4°C and moved to a Percival growth chamber maintained at 21°C with constant light. *S. irio* seeds were stratified for 10 days in the dark at 4°C and moved to a Percival or Phytotron growth chamber maintained at 21°C with constant light. 10–14-day-old seedlings were transplanted to soil and grown under long-day conditions: 16 hours of light at 21°C and 8 hours of dark at 18°C.

In the case of *A. lyrata*, sterilized seeds were suspended in sterile deionized water and placed in the dark at 4°C for 4 weeks. After 4 weeks, *A. lyrata* seeds were dried and plated on MS plates with 2% sucrose, stratified for 10 days in the dark at 4°C, and moved to a Percival growth chamber that was maintained at 21°C with constant light. 10–14-day-old seedlings were then transplanted to soil and grown under long-day conditions: 16 hours of light at 20°C and 8 hours of dark at 16°C. Plants were kept in these conditions for 2 months before vernalization at 4°C with 20 hours of light and 4 hours of dark for 9 weeks. After vernalization, plants were moved back to the growth chamber with 16 hours of light at 21°C and 8 hours of dark at 16°C.

All *A. thaliana* seeds used in this study are the *Columbia* (*Col-0*) ecotype. The following constructs were transformed into the homozygous *lre-7/lre-7* mutant background:*AtLRE_pro_:AtLRE-cYFP, AtLRE_pro_:AtLRE(E47P)-cYFP, AtLRE_pro_:AtLRE(R120G)-cYFP, AtLRE_pro_:AtLRE(R120T)-cYFP, AtLRE_pro_:AtLRE(E47P-R120G)-cYFP, AtLRE_pro_:AlyLRE-cYFP,* and *AtLRE_pro_:SiLRE-cYFP.* Transgenic plants were selected on plates containing both hygromycin B and glufosinate ammonium.

### Cloning transgenic constructs

The *AtLRE_pro_:AtLRE(E47P)-cYFP, AtLRE_pro_:AtLRE(R120G)-cYFP, AtLRE_pro_:AtLRE(E47P-R120G)-cYFP,* and *AtLRE_pro_:AtLRE (R120T)-cYFP* transgenic lines were created by introducing nucleotide changes that will cause E to P, R to G, and R to T changes, respectively, in the LRE protein sequence in the previously published construct: *AtLRE_pro_:LRE-cYFP* (Liu et al. 2016). The *AtLRE_pro_:AlyLRE-cYFP* and *AtLRE_pro_:SiLRE-cYFP* transgenic lines were created following the protocol established with the previously published *AtLRE_pro_:LRE-cYFP* construct (Liu et al. 2016). Mutations were introduced by PCR with PrimeSTAR® GXL DNA Polymerase (TaKaRa Bio Inc.; Catalog # R050A) using primers and DNA templates listed in Table S6. Overlap PCR was performed for *AtLRE_pro_:AtLRE(E47P-R120G)-cYFP*, *AtLRE_pro_:AlyLRE-cYFP* and *AtLRE_pro_:SiLRE-cYFP* as indicated in Table S6prior to cloning. The amplified PCR products were cloned into *AtLRE_pro_:LRE-cYFP* plasmid linearized with *SpeI-HF* (NEB, Catalog # R3133S) and *AscI* (NEB, Catalog # R0558S) by using the In-Fusion HD Cloning Plus system (Clontech, Catalog # 639645). The recombinant plasmids were transformed into Stellar™ Competent Cells (Clontech, Catalog # 636763), and positive colonies were selected on LB plates containing Spectinomycin (100 µg/mL, Sigma-Aldrich, Catalog # 85555).

All constructs generated were sequence-verified for the introduced point mutations (Eton Bioscience, Inc.) before transforming into *Agrobacterium tumefaciens* (GV3101 pMP90 strain). The positive colony selected for transformation into *A. thaliana* was also verified by colony PCR for the presence of the transgene.

### Plant transformation

Transformation solution containing *Agrobacterium tumefaciens* (GV3101 pMP90 strain) harboring the desired transgene was sprayed onto *A. thaliana* inflorescences (Chung et al. 2000). Hygromycin-resistant T1 transformants were selected as described (Harrison et al. 2006). Briefly, T1 seeds were plated and stratified in the dark at 4°C for 2-3 days. Plates were then placed in a Percival growth chamber set at 21°C with continuous light (70-100 µmol·m^-2^·s^-1^) for 5-6 hours, then placed in the dark at room temperature for 3-4 days. Plates were then returned to the Percival growth chamber, and transformants were selected based on the presence of true leaves, which were present only in hygromycin-resistant plants.

### Identification of single-loci insertion lines

For each construct, at least 15 T1 hygromycin-resistant transformants were transplanted to soil. Among these lines, candidate single insertion lines were identified based on segregation of resistance to hygromycin B among T2 progeny. T1 plants are expected to be heterozygous for the transgene at the insertion locus. T1 plants that gave rise to progeny with a 3:1 ratio of resistant to sensitive plants were considered as candidate single locus insertion lines. 15 T2 plants of candidate lines were transplanted to soil, and T3 selfed seeds were collected to identify homozygous lines in T3 populations. Those T2 plants that gave rise to T3 progeny that segregated 100% resistance to hygromycin B were homozygous for the transgene.

### Genomic DNA isolation from *A. thaliana* seeds

Genomic DNA was extracted from dry *A. thaliana* seeds (50 mg) using a CTAB-based method adapted from Dellaporta et al. (1983). Seeds were frozen in liquid nitrogen, ground to a fine powder, and suspended in CTAB B solution (1.4 M NaCl, 0.1 M Tris–HCl (pH 8.0), 2% (w/v) CTAB). After incubation at 65°C, the homogenate was treated with a 1:1 mixture of CTAB B and CTAB C (50 mM Tris–HCl (pH 8.0), 10 mM EDTA (pH 8.0), 1% (w/v) CTAB) and centrifuged to remove debris. The supernatant was extracted twice with chloroform solution (chloroform: isoamyl alcohol, 24:1), followed by precipitation with CTAB C. The resulting DNA pellet was resuspended in CTAB D (50 mM Tris–HCl (pH 8.0), 10 mM EDTA (pH 8.0), 1 M CaCl, 1% (w/v) CTAB) containing RNase A and incubated at 37°C. DNA was subsequently precipitated with ethanol, washed with 75% ethanol, and resuspended in TE buffer. DNA yield and purity were assessed spectrophotometrically at 260 nm, typically ranging between 200 ng/mL and 1 mg/mL, and these samples were subsequently used to determine transgene copy number using digital droplet PCR.

### Digital droplet PCR

Genomic DNA from selected transgenic lines that were designated as single-locus insertion lines was isolated and tested to check whether they had any variation in the copy number of the transgene inserted in that locus using digital droplet PCR (ddPCR) (Figure S3) as follows: ddPCR was performed using QX200TM ddPCR EvaGreen Supermix on a Bio-Rad QX200 Droplet Digital PCR system. The single-copy gene PMR4 was used as a reference (Dong et al. 2015). Two sets of specific oligonucleotides were designed to target the PMR4 gene or the cYFP coding sequence. For the ddPCR reaction mix, a 20 µL ddPCR supermix was made for each gene using 10ng template genomic DNA and 50nm oligos mix. Each reaction was loaded into 8-well cartridges for droplet generation and then transferred to the QX200 Droplet Digital PCR system, with a standard ddPCR reaction step of 10 minutes at 95°C, 40 cycles of 94°C for 30 seconds and an annealing temperature of 58 °C for 60 seconds, and a final 10 minutes at 98°C. The ddPCR data were analyzed using QuantaSoft analysis (Bio-Rad) and plotted using gPlot database.

### Citrine YFP expression in ovules

Expression of the cYFP in the FA of synergid cells was examined and imaged with a Zeiss LSM880 inverted confocal microscope. For cYFP imaging, samples were excited with a 488-nm laser line, and emission was collected between 510 and 550 nm. YFP images were processed with ZEN Black and ImageJ software (http://imagej.nih.gov/ij/; Schindelin et al. 2012). ImageJ was used to insert scale bars and assemble the final images.

### Standardization of inter-lineage crosses

Plants that were 4 or 5 weeks old after they were transplanted to soil were used as male and female parents. Stage 12c buds were emasculated as previously described (Smyth et al. 1990). Twenty-four hours after emasculation, pistils were manually and profusely pollinated with *A. thaliana, A. lyrata,* or *S. irio* pollen. Pistils were collected 24 hours after pollination for aniline blue staining.

### Aniline blue staining

Aniline blue staining was performed as described (Mori et al. 2006; Noble and Palanivelu 2020). Pistils were stained for 24 hours and mounted in decolorized aniline blue solution (DABS) containing 50% glycerol, then sealed with clear nail polish (Sally Hansen, Hard as Nails). Pollen tube-ovule interactions were scored under a Zeiss Axiovert 100 fluorescent microscope. Images were captured using Metamorph (Version 7; Molecular Devices) with an IR filter and auto-exposure settings. The number of pistils and ovules scored is indicated in each figure.

### Classification and quantification of pollen tube reception outcomes

Pollen tube behavior in aniline blue-stained pistils was classified into four categories. Type 1 comprised ovules for which no pollen tube was observed at the micropyle. Type 2 comprised ovules in which a pollen tube reached the micropyle but did not enter the female gametophyte. Type 3 comprised ovules in which a pollen tube entered the micropyle but continued to grow and coil within the female gametophyte, providing morphological evidence of failed pollen tube reception. Type 4 comprised ovules in which a pollen tube entered the female gametophyte and terminated near the expected position of the synergid cells without visible continued growth or coiling.

All four categories were scored in each cross. For comparisons of PTR after pollen tube entry, failure was calculated as the percentage of pollen tube-entered ovules exhibiting overgrowth [%OG/PEO = Type 3/(Type 3 + Type 4) × 100]. Types 1 and 2 were excluded from this calculation because the pollen tubes had not entered the female gametophyte and PTR therefore could not be assessed. Consequently, %OG/PEO measures PTR failure conditional on pollen tube entry rather than the total reproductive barrier, which may also include defects occurring before entry.

Classification as Type 3 required unequivocal evidence of continued pollen tube growth or coiling within the female gametophyte. Type 4 classification was based on the lack of apparent overgrowth and did not directly establish pollen tube rupture, sperm release, or fertilization. Some Type 4 outcomes could therefore represent pollen tube arrest without productive reception. Accordingly, %OG/PEO should be interpreted as a conservative estimate of morphologically evident PTR failure among pollen tube-entered ovules.

### Statistical analysis

For each crossed pistil, the proportion of ovules that received an abnormal pollen tube was calculated by dividing the number of ovules with abnormal pollen tube behavior (type 3) by the total number of ovules that received a pollen tube (both abnormal (type 3) and normal (type 4); Fig. 1B to F). This proportion was then subjected to an arcsine square root transformation using Microsoft Excel. Transformed values were used in two-sample *t*-tests comparing each test line to the *AtLRE_pro_:AtLRE-cYFP* control line, except in Fig. 1G, where values were compared to wild-type *A. thaliana* values, with a significance level of 0.05. Significant values are indicated in the graphs as follows: where ns > p= 0.05, *p < 0.05, **p < 0.01, ***p < 0.001, ****p < 0.0001. Graphs show the average percentage of pollen tube overgrowth per pollen tube-entered ovule (OG/PEO) across all siliques with pollen tube reception for each cross. Data were plotted using Prism 10 (https://www.graphpad.com/).

Control genotypes (Col-0, *lre-7*, and *lre-7* complemented with *AtLRE* (*AtLRE*pro:*AtLRE*-cYFP)) were included in each experimental series; therefore, control values reported in each figure/graph represent the contemporaneous controls measured in the same experimental block as the corresponding test lines, rather than values pooled across experiments.

### Identifying orthologs of *LRE* and *LLG1*

Genomes were obtained for *Arabidopsis thaliana,Brassica rapa*, *Carica papaya*, *Capsella rubella*, *Camelina sativa*, *Cleome violacea*, *Descurainia sophioides*, *Eutrema salsugineum*, *Euclidium syriacum*, *Gossypium raimondii*, *Lunaria annua*, *Lepidium sativum*, *Malcolmia maritima*, *Myagrum perfoliatum*, *Rorippa islandica*, *Sinapis alba*, *Sisymbrium irio*, *Schrenkiella parvula*, *Stanleya pinnata*, *Thlaspi arvense*, and *Theobroma cacao* from Phytozome, v13 (Goodstein et al. 2012); *Aethionema arabicum* from *Ae. arabicum* DB (Fernandez-Pozo et al. 2021); and for *Tarrenaya hassleriana* (Cheng et al. 2013), *Bretschneidera sinensis* (Liu et al. 2022), *Moringa oleifera* (Chang et al. 2022), *Capparis spinosa* (Wang et al. 2022), and *Tropaeolum majus* (Friedhoff et al. 2024) from their respective publications (Table S1). If available, coding sequence (CDS) genomes were obtained. For *B. sinensis* and *T. hassleriana*, whole-genome and annotation files were used as input into GFFRead, v0.12.7, to generate the CDS genome, using the command: gffread -g genome.fa annotation.gff3 -x cds.fa (Pertea and Pertea 2020). Alternative transcripts were removed using primary_transcript.py from Orthofinder, v2.5.5 (Emms and Kelly 2015; Emms and Kelly 2019). CDS genomes were translated into peptide genomes using SeqKit2, v2.9.0, with the command: seqkit translate cds.fa > pep.fa (Shen et al. 2016; Shen et al. 2024). Using previously published phylogenies of Brassicaceae and Brassicales for reference (Hall et al. 2004; Beilstein et al. 2006; Beilstein et al. 2008; Edger et al. 2018; Hendriks et al. 2023), a Newick tree, newick.txt, was made for the selected species within Mesquite, v3.81 (Maddison and Maddison 2023). Homologs of the selected genomes were identified and assigned to Orthogroups using Orthofinder, v2.5.5, with the command: orthofinder -f /path/to/peptide_genomes -s newick.txt -y -X -M msa -t 64 (Emms and Kelly, 2015; Emms and Kelly, 2019). *LRE* and *LLG1* homologues were found in two Orthogroups: one containing all Brassicaceae *LRE* orthologs and the other containing all Brassicaceae *LLG1* orthologs and non-Brassicaceae *LRE/LLG1* homologues. Together, these two Orthogroups contained all previously identified Brassicales and Malvales *LRE/LLG1* homologues from Noble et al. 2022 in the species selected (Table S2).

### *LRE* and *LLG1* CDS alignments and phylogenies

*LRE* and *LLG1* peptide sequences were aligned using the PPP algorithm in MUSCLE, v5.2.linux64, using the command: muscle -align pep_sequences.fa -output pep_alignment.afa -threads 20 (Edgar 2022). The peptide alignment was reverse-translated into a nucleotide alignment using PAL2NAL, v14, with the command: pal2nal.pl pep_alignment.afa nuc_sequences.fa -output fasta > nuc_alignment.afa (Suyama et al. 2006). The nucleotide alignment was then trimmed at the codon level with the minimum block length set to 5 and allowing half of the gap positions and otherwise default parameters using Gblocks, v0.91b, yielding trimmed_nuc_alignment.afa (Talavera and Castresana 2007). Phylogeny of the *LRE* and *LLG1* homologues was then inferred in RAxML-ng, v1.2.2-master, starting with 10 random trees and 10 parsimony trees using the GTR+G model of evolution and 1000 Felsenstein bootstrap replicates with the command: raxml-ng -all -msa trimmed_nuc_alignment.afa -model GTR+G -bs-trees 1000 (Kozlov et al. 2019). The best tree was then visualized using iTOL, v7 (Letunic and Bork 2024) and rooted by the Malvales *LRE/LLG1* clade, which served as an outgroup.

### PAML Analysis

PAML analyses were performed with PAML, v4.10.9 (Yang 2007), using the alignment, trimmed_nuc_alignment.afa, and best tree from RAxML-ng rooted by the Malvales *LRE/LLG1* clade, as described above. The guidance of Álvarez-Carretero et al. 2023 was followed for running the site, branch, and branch-site models. Alternative models were compared against their corresponding null model using the likelihood-ratio test (LRT). To test for multiple site classes, M1a (Nearly Neutral) was compared against M0 (one-ratio) (Goldman and Yang 1994; Nielsen and Yang 1998; Wong et al. 2004; Yang et al. 2005). The branch tests were performed four times, with different branches highlighted as the foreground: the post-duplication branch (PDB) or all clade branches, including the PDB, in either *LRE* or *LLG1* were highlighted as the foreground. For the branch tests, the alternative models were compared against M0 (Yang 1998; Yang and Nielsen 1998). For the Model C clade tests, all branches within the *LRE* clade, including the PDB, were highlighted as foreground, and the alternative model was compared against M2a_rel (Bielawski and Yang 2004; Weadick and Chang 2012). The Model A branch-sites test was performed on either the PDB or the entire clade, including the PDB, for *LRE*. For the Model A branch-site tests, the alternative model, where ω was set as a free parameter starting at a value of 0.5, was compared against the null model, where ω was fixed at a value of 1 (Yang et al. 2000; Yang and Nielsen 2002; Yang et al. 2005; Zhang et al. 2005). Posterior probabilities for site classes were determined using Bayes empirical Bayes (BEB) analysis (Yang et al. 2005). Results from PAML, including parameter estimates, likelihood scores, likelihood-ratio tests, and BEB results for Model A branch-site tests, are recorded in Table S5.

### Structural Modeling and Analysis

AlphaFold2 was used with 32 seeds to generate 160 models for analysis consistent with established protocols (Monteiro Da Silva et al. 2024; Galeazzi et al. 2025). Multiple Sequence Alignments for each variant were generated using the JackHMMR method and respective sequences as queries (Finn et al. 2011). After various subsampling conditions were tested by modifying the *max_seq* and *extra_seq* for WT and each AtLRE variant, a *max_seq*:*extra_seq* of 128:256 was selected for downstream analyses. Any structures with pTM <50 and pLDDT <70 were filtered out in accordance with established protocols (Jumper et al. 2021; Abramson et al. 2024).

The predictions were then loaded as trajectories into MDAnalysis in Python 3.12.3 (Michaud-Agrawal et al. 2011; Gowers et al. 2016). Structures were first superimposed and aligned to the conserved core region using the AlignTraj function in MDAnalysis. Root mean square deviation (RMSD) with an all-atom Cα selection was calculated relative to the top-ranked structure for each variant. Plots were generated using Matplotlib (Hunter 2007). Visual rendering of proteins was generated using PyMOL (version 2.3.0).

## Acknowledgments

E.J. and M.J.A. were supported by funding from the U.S. National Science Foundation awarded to RP (IOS-1146090) and MAB (DBI-2019674), respectively, and by the School of Plant Sciences. In addition, M.J.A. received funding from the U.S. National Institutes of Health Initiative to Maximize Student Development (T32GM139779). J.A.N. was supported by the following: IGERT Comparative Genomics Program at the University of Arizona (Award ID: 0654435); NSF Graduate Research Fellowship: Grant DGE-1143953; the Boynton Graduate Fellowship; and the University of Arizona Graduate College Office of Diversity and Inclusion. We thank Dr. Zuzana Gadiou and Dr. Vinod Kumar for assistance with interspecific crosses done at the Institute of Experimental Botany of the Czech Academy of Sciences. We thank Dr. Dean Billheimer, Zuckerman College of Public Health, University of Arizona, for advising us on statistical analysis of the data presented in this study. We thank Dr. Mark Johnson, Dr. Ueli Grossniklaus, Dr. Steven Smith, Dr. Ramin Yadegari, and Dr. Rebecca Mosher for helpful discussions.

## References

Abramson, J., Adler, J., Dunger, J., Evans, R., Green, T., Pritzel, A., Ronneberger, O., Willmore, L., Ballard, A. J., Bambrick, J., Bodenstein, S. W., Evans, D. A., Hung, C.-C., O’Neill, M., Reiman, D., Tunyasuvunakool, K., Wu, Z., Žemgulytė, A., Arvaniti, E., … Jumper, J. M. (2024). Accurate structure prediction of biomolecular interactions with AlphaFold 3. Nature, 630(8016), 493–500. 10.1038/s41586-024-07487-w

Álvarez-Carretero, S., Kapli, P., & Yang, Z. (2023). Beginner’s Guide on the Use of PAML to Detect Positive Selection. Molecular Biology and Evolution, 40(4), msad041. 10.1093/molbev/msad041

Amien, S., Kliwer, I., Márton, M. L., Debener, T., Geiger, D., et al. (2010). Defensin-like ZmES4 mediates pollen tube burst in maize via opening of the potassium channel KZM1. PLOS Biol. 8:e100038810.1371/journal.pbio.1000388

Baack, E., Melo, M. C., Rieseberg, L. H., & Ortiz-Barrientos, D. (2015). The origins of reproductive isolation in plants. New Phytologist, 207(4), 968–984. 10.1111/nph.13424

Beilstein, M. A., Al-Shehbaz, I. A., & Kellogg, E. A. (2006). Brassicaceae phylogeny and trichome evolution. American Journal of Botany, 93(4), 607–619. 10.3732/ajb.93.4.607

Beilstein, M. A., Al-Shehbaz, I. A., Mathews, S., & Kellogg, E. A. (2008). Brassicaceae phylogeny inferred from phytochrome A and *ndhF* sequence data: Tribes and trichomes revisited. American Journal of Botany, 95(10), 1307–1327. 10.3732/ajb.0800065

Bielawski, JosephP., & Yang, Z. (2004). A Maximum Likelihood Method for Detecting Functional Divergence at Individual Codon Sites, with Application to Gene Family Evolution. Journal of Molecular Evolution, 59(1). 10.1007/s00239-004-2597-8

Cao, Y., Cui, X., Yang, Y., Pan, L., Yang, F., Li, S., Wu, D., Ding, Y., Chen, R., Wang, N., et al. (2025). Pan-family pollen signals control an interspecific stigma barrier across Brassicaceae species. Science 391, Article eady2347. 10.1126/science.ady2347

Capron, A., Gourgues, M., Neiva, L. S., Faure, J.-E., Berger, F., Pagnussat, G., Krishnan, A., Alvarez-Mejia, C., Vielle-Calzada, J.-P., Lee, Y.-R., Liu, B., & Sundaresan, V. (2008). Maternal Control of Male-Gamete Delivery in *Arabidopsis* Involves a Putative GPI-Anchored Protein Encoded by the *LORELEI* Gene. The Plant Cell, 20(11), 3038– 3049. 10.1105/tpc.108.061713

Chang, J., Marczuk-Rojas, J. P., Waterman, C., Garcia-Llanos, A., Chen, S., Ma, X., Hulse-Kemp, A., Van Deynze, A., Van De Peer, Y., & Carretero-Paulet, L. (2022). Chromosome-scale assembly of the *Moringa oleifera* Lam. Genome uncovers polyploid history and evolution of secondary metabolism pathways through tandem duplication. The Plant Genome, 15(3), e20238. 10.1002/tpg2.20238

Cheng, S., Van Den Bergh, E., Zeng, P., Zhong, X., Xu, J., Liu, X., Hofberger, J., De Bruijn, S., Bhide, A. S., Kuelahoglu, C., Bian, C., Chen, J., Fan, G., Kaufmann, K., Hall, J. C., Becker, A., Bräutigam, A., Weber, A. P. M., Shi, C., … Schranz, M. E. (2013). The *Tarenaya hassleriana* Genome Provides Insight into Reproductive Trait and Genome Evolution of Crucifers. The Plant Cell, 25(8), 2813–2830. 10.1105/tpc.113.113480

Chung, M.-H., Chen, M.-K., & Pan, S.-M. (2000). Floral Spray Transformation Can Efficiently Generate Arabidopsis. Transgenic Research, 9(6), 471–486. 10.1023/A:1026522104478

Dellaporta, S. L., Wood, J., & Hicks, J. B. (1983). A plant DNA minipreparation: Version II. Plant Molecular Biology Reporter, 1(4), 19–21. 10.1007/BF02712670

Dong, L., Meng, Y., Sui, Z., Wang, J., Wu, L., & Fu, B. (2015). Comparison of four digital PCR platforms for accurate quantification of DNA copy number of a certified plasmid DNA reference material. Scientific Reports, 5(1), 13174. 10.1038/srep13174

Edgar, R. C. (2022). Muscle5: High-accuracy alignment ensembles enable unbiased assessments of sequence homology and phylogeny. Nature Communications, 13(1), 6968. 10.1038/s41467-022-34630-w

Edger, P. P., Hall, J. C., Harkess, A., Tang, M., Coombs, J., Mohammadin, S., Schranz, M. E., Xiong, Z., Leebens-Mack, J., Meyers, B. C., Sytsma, K. J., Koch, M. A., Al-Shehbaz, I. A., & Pires, J. C. (2018). Brassicales phylogeny inferred from 72 plastid genes: A reanalysis of the phylogenetic localization of two paleopolyploid events and origin of novel chemical defenses. American Journal of Botany, 105(3), 463–469. 10.1002/ajb2.1040

Emms, D. M., & Kelly, S. (2015). OrthoFinder: Solving fundamental biases in whole genome comparisons dramatically improves orthogroup inference accuracy. Genome Biology, 16(1), 157. 10.1186/s13059-015-0721-2

Emms, D. M., & Kelly, S. (2019). OrthoFinder: Phylogenetic orthology inference for comparative genomics. Genome Biology, 20(1), 238. 10.1186/s13059-019-1832-y

Escobar-Restrepo, J.-M., Huck, N., Kessler, S., Gagliardini, V., Gheyselinck, J., Yang, W.-C., & Grossniklaus, U. (2007). The FERONIA Receptor-like Kinase Mediates Male-Female Interactions During Pollen Tube Reception. Science, 317(5838), 656–660. 10.1126/science.1143562

Feng, H., Liu, C., Fu, R., Zhang, M., Li, H., Shen, L., Wei, Q., Sun, X., Xu, L., Ni, B., & Li, C. (2019). LORELEI-LIKE GPI-ANCHORED PROTEINS 2/3 Regulate Pollen Tube Growth as Chaperones and Coreceptors for ANXUR/BUPS Receptor Kinases in Arabidopsis. Molecular Plant, 12(12), 1612–1623. 10.1016/j.molp.2019.09.004

Fernandez-Pozo, N., Metz, T., Chandler, J. O., Gramzow, L., Mérai, Z., Maumus, F., Mittelsten Scheid, O., Theißen, G., Schranz, M. E., Leubner-Metzger, G., & Rensing, S. A. (2021). Aethionema arabicum genome annotation using PacBio full-length transcripts provides a valuable resource for seed dormancy and Brassicaceae evolution research. The Plant Journal: For Cell and Molecular Biology, 106(1), 275–293. 10.1111/tpj.15161

Finn, R. D., Clements, J., & Eddy, S. R. (2011). HMMER web server: Interactive sequence similarity searching. Nucleic Acids Research, 39(suppl), W29–W37. 10.1093/nar/gkr367

Fleming, J., Magana, P., Nair, S., Tsenkov, M., Bertoni, D., Pidruchna, I., Lima Afonso, M. Q., Midlik, A., Paramval, U., Žídek, A., Laydon, A., Kovalevskiy, O., Pan, J., Cheng, J., Avsec, Ž., Bycroft, C., Wong, L. H., Last, M., Mirdita, M., … Velankar, S. (2025). AlphaFold Protein Structure Database and 3D-Beacons: New Data and Capabilities. Journal of Molecular Biology, 437(15), 168967. 10.1016/j.jmb.2025.168967

Friedhoff, R., Wittstock, U., & Pucker, B. (2024). *Genome sequence of the medicinal plant* Tropaeolum majus *provides insights into flavonoid biosynthesis*. Plant Biology. 10.1101/2024.11.01.621629

Fujii, S., Tsuchimatsu, T., Kimura, Y., Ishida, S., Tangpranomkorn, S., Shimosato-Asano, H., Iwano, M., Furukawa, S., Itoyama, W., Wada, Y., Shimizu, K. K., & Takayama, S. (2019). A stigmatic gene confers interspecies incompatibility in the Brassicaceae. Nature Plants, 5(7), 731–741. 10.1038/s41477-019-0444-6

Fujii, S., Yamamoto, E., Ito, S., Tangpranomkorn, S., Kimura, Y., et al. (2023). SHI family transcription factors regulate an interspecific barrier. Nat. Plants 9, 1862–73. 10.1038/s41477-023-01535-5

Galeazzi, F. M., Monteiro Da Silva, G., Arantes, P., Varghese, I., Shukla, A., & Rubenstein, B. M. (2025). FastConformation: A Standalone ML-Based Toolkit for Modeling and Analyzing Protein Conformational Ensembles at Scale. Biophysics. 10.1101/2025.05.09.653048

Galindo-Trigo, S., Blanco-Touriñán, N., DeFalco, T. A., Wells, E. S., Gray, J. E., Zipfel, C., & Smith, L. M. (2020). *Cr*RLK1L receptor-like kinases HERK1 and ANJEA are female determinants of pollen tube reception. EMBO Reports, 21(2). 10.15252/embr.201948466

Gao, Q., Wang, C., Xi, Y., Shao, Q., Li, L., & Luan, S. (2022). A receptor–channel trio conducts Ca2+ signalling for pollen tube reception. Nature, 607(7919), 534–539. 10.1038/s41586-022-04923-7

Ge, Z., Bergonci, T., Zhao, Y., Zou, Y., Du, S., Liu, M.-C., Luo, X., Ruan, H., García-Valencia, L. E., Zhong, S., Hou, S., Huang, Q., Lai, L., Moura, D. S., Gu, H., Dong, J., Wu, H.-M., Dresselhaus, T., Xiao, J., … Qu, L.-J. (2017). *Arabidopsis* pollen tube integrity and sperm release are regulated by RALF-mediated signaling. Science, 358(6370), 1596–1600. 10.1126/science.aao3642

Ge, Z., Zhao, Y., Liu, M.-C., Zhou, L.-Z., Wang, L., Zhong, S., Hou, S., Jiang, J., Liu, T., Huang, Q., Xiao, J., Gu, H., Wu, H.-M., Dong, J., Dresselhaus, T., Cheung, A. Y., & Qu, L.-J. (2019). LLG2/3 Are Co-receptors in BUPS/ANX-RALF Signaling to Regulate Arabidopsis Pollen Tube Integrity. Current Biology, 29(19), 3256–3265.e5. 10.1016/j.cub.2019.08.032

Goldman, N., & Yang, Z. (1994). A codon-based model of nucleotide substitution for protein-coding DNA sequences. Molecular Biology and Evolution. 10.1093/oxfordjournals.molbev.a040153

Goodstein, D. M., Shu, S., Howson, R., Neupane, R., Hayes, R. D., Fazo, J., Mitros, T., Dirks, W., Hellsten, U., Putnam, N., & Rokhsar, D. S. (2012). Phytozome: A comparative platform for green plant genomics. Nucleic Acids Research, 40(D1), D1178–D1186. 10.1093/nar/gkr944

Gowers, R., Linke, M., Barnoud, J., Reddy, T., Melo, M., Seyler, S., Domański, J., Dotson, D., Buchoux, S., Kenney, I., & Beckstein, O. (2016). MDAnalysis: A Python Package for the Rapid Analysis of Molecular Dynamics Simulations. 98–105. 10.25080/Majora-629e541a-00e

Hall, J. C., Iltis, H. H., & Sytsma, K. J. (2004). Molecular Phylogenetics of Core Brassicales, Placement of Orphan Genera Emblingia, Forchhammeria, Tirania, and Character Evolution. Systematic Botany, 29(3), 654–669. JSTOR.

Harrison, S. J., Mott, E. K., Parsley, K., Aspinall, S., Gray, J. C., & Cottage, A. (2006). A rapid and robust method of identifying transformed Arabidopsis thaliana seedlings following floral dip transformation. Plant Methods, 2(1), 19. 10.1186/1746-4811-2-19

Hendriks, K. P., Kiefer, C., Al-Shehbaz, I. A., Bailey, C. D., Hooft Van Huysduynen, A., Nikolov, L. A., Nauheimer, L., Zuntini, A. R., German, D. A., Franzke, A., Koch, M. A., Lysak, M. A., Toro-Núñez, Ó., Özüdoğru, B., Invernón, V. R., Walden, N., Maurin, O., Hay, N. M., Shushkov, P., … Lens, F. (2023). Global Brassicaceae phylogeny based on filtering of 1,000-gene dataset. Current Biology, 33(19), 4052–4068.e6. 10.1016/j.cub.2023.08.026

Higashiyama, T., Inatsugi, R., Sakamoto, S., Sasaki, N., Mori, T., Kuroiwa, H., Nakada, T., Nozaki, H., Kuroiwa, T., & Nakano, A. (2006). Species Preferentiality of the Pollen Tube Attractant Derived from the Synergid Cell of *Torenia fournieri*. Plant Physiology, 142(2), 481–491. 10.1104/pp.106.083832

Hou, Y., Guo, X., Cyprys, P., Zhang, Y., Bleckmann, A., et al. (2016). Maternal ENODLs are required for pollen tube reception in Arabidopsis. Curr. Biol. 26:2343–50

Huang, J., Yang, L., Yang, L., Wu, X., Cui, X., Zhang, L., Hui, J., Zhao, Y., Yang, H., Liu, S., Xu, Q., Pang, M., Guo, X., Cao, Y., Chen, Y., Ren, X., Lv, J., Yu, J., Ding, J., … Duan, Q. (2023). Stigma receptors control intraspecies and interspecies barriers in Brassicaceae. Nature, 614(7947), 303–308. 10.1038/s41586-022-05640-x

Hunter, J. D. (2007). Matplotlib: A 2D Graphics Environment. Computing in Science & Engineering, 9(3), 90–95. 10.1109/MCSE.2007.55

Ju, Y., Yuan, J., Jones, D. S., Zhang, W., Staiger, C. J., & Kessler, S. A. (2021). Polarized NORTIA accumulation in response to pollen tube arrival at synergids promotes fertilization. Developmental Cell, 56(21), 2938–2951.e6. 10.1016/j.devcel.2021.09.026

Jumper, J., Evans, R., Pritzel, A., Green, T., Figurnov, M., Ronneberger, O., Tunyasuvunakool, K., Bates, R., Žídek, A., Potapenko, A., Bridgland, A., Meyer, C., Kohl, S. A. A., Ballard, A. J., Cowie, A., Romera-Paredes, B., Nikolov, S., Jain, R., Adler, J., … Hassabis, D. (2021). Highly accurate protein structure prediction with AlphaFold. Nature, 596(7873), 583–589. 10.1038/s41586-021-03819-2

Kozlov, A. M., Darriba, D., Flouri, T., Morel, B., & Stamatakis, A. (2019). RAxML-NG: A fast, scalable and user-friendly tool for maximum likelihood phylogenetic inference. Bioinformatics, 35(21), 4453–4455. 10.1093/bioinformatics/btz305

Lan, Z., Song, Z., Wang, Z., Li, L., Liu, Y., Zhi, S., Wang, R., Wang, J., Li, Q., Bleckmann, A., Zhang, L., Dresselhaus, T., Dong, J., Gu, H., Zhong, S., & Qu, L.-J. (2023). Antagonistic RALF peptides control an intergeneric hybridization barrier on Brassicaceae stigmas. Cell, 186(22), 4773–4787.e12. 10.1016/j.cell.2023.09.003

Letunic, I., & Bork, P. (2024). Interactive Tree of Life (iTOL) v6: Recent updates to the phylogenetic tree display and annotation tool. Nucleic Acids Research, 52(W1), W78– W82. 10.1093/nar/gkae268

Leydon, A. R., Tsukamoto, T., Dunatunga, D., Qin, Y., Johnson, M. A., Palanivelu, R. (2015). Pollen tube discharge completes the process of synergid degeneration that is initiated by pollen tube–synergid interaction in Arabidopsis. Plant Physiol. 169:485–96. 10.1104/pp.15.00528

Leydon, A. R., Beale, K. M., Woroniecka, K., Castner, E., Chen, J., Horgan, C., Palanivelu, R., & Johnson, M. A. (2013). Three MYB transcription factors control pollen tube differentiation required for sperm release. Current Biology: CB, 23(13), 1209–1214. 10.1016/j.cub.2013.05.021

Liu, C., Shen, L., Xiao, Y., Vyshedsky, D., Peng, C., Sun, X., Liu, Z., Cheng, L., Zhang, H., Han, Z., Chai, J., Wu, H.-M., Cheung, A. Y., & Li, C. (2021). Pollen PCP-B peptides unlock a stigma peptide–receptor kinase gating mechanism for pollination. Science, 372(6538), 171–175. 10.1126/science.abc6107

Liu, H.-L., Harris, A. J., Wang, Z.-F., Chen, H.-F., Li, Z.-A., & Wei, X. (2022). The genome of the Paleogene relic tree *Bretschneidera sinensis*: Insights into trade-offs in gene family evolution, demographic history, and adaptive SNPs. DNA Research, 29(1), dsac003. 10.1093/dnares/dsac003

Liu, X., Castro, C., Wang, Y., Noble, J., Ponvert, N., Bundy, M., Hoel, C., Shpak, E., & Palanivelu, R. (2016). The Role of LORELEI in Pollen Tube Reception at the Interface of the Synergid Cell and Pollen Tube Requires the Modified Eight-Cysteine Motif and the Receptor-Like Kinase FERONIA. The Plant Cell, 28(5), 1035–1052. 10.1105/tpc.15.00703

Liu, Z., Chu, X., Ren, W., Cheng, L., Liu, C., Wang, C., Gao, S., Dai, S., Li, C. 2025. PCP-B peptides and CrRLK1L receptor kinases control pollination via pH gating of aquaporins in Arabidopsis. Developmental Cell, 60, 1336–1347. 10.1016/j.devcel.2024.12.026

Maddison, W. P., & Maddison, D. R. (n.d.). Mesquite: A modular system for evolutionary analysis (Version 3.812023) [Computer software]. Retrieved http://www.mesquiteproject.org

Márton, M. L., Fastner, A., Uebler, S., & Dresselhaus, T. (2012). Overcoming Hybridization Barriers by the Secretion of the Maize Pollen Tube Attractant ZmEA1 from Arabidopsis Ovules. Current Biology, 22(13), 1194–1198. 10.1016/j.cub.2012.04.061

Michaud-Agrawal, N., Denning, E. J., Woolf, T. B., & Beckstein, O. (2011). MDAnalysis: A toolkit for the analysis of molecular dynamics simulations. Journal of Computational Chemistry, 32(10), 2319–2327. 10.1002/jcc.21787

Monteiro Da Silva, G., Cui, J. Y., Dalgarno, D. C., Lisi, G. P., & Rubenstein, B. M. (2024). High-throughput prediction of protein conformational distributions with subsampled AlphaFold2. Nature Communications, 15(1), 2464. 10.1038/s41467-024-46715-9

Mori, T., Kuroiwa, H., Higashiyama, T., & Kuroiwa, T. (2006). GENERATIVE CELL SPECIFIC 1 is essential for angiosperm fertilization. Nature Cell Biology, 8(1), 64–71. 10.1038/ncb1345

Müller, L. M., Lindner, H., Pires, N. D., Gagliardini, V., & Grossniklaus, U. (2016). A subunit of the oligosaccharyltransferase complex is required for interspecific gametophyte recognition in Arabidopsis. Nature Communications, 7(1), 10826. 10.1038/ncomms10826

Nielsen, R., & Yang, Z. (1998). Likelihood Models for Detecting Positively Selected Amino Acid Sites and Applications to the HIV-1 Envelope Gene. Genetics, 148(3), 929– 936. 10.1093/genetics/148.3.929

Noble, J. A., Bielski, N. V., Liu, M.-C. J., DeFalco, T. A., Stegmann, M., Nelson, A. D. L., McNamara, K., Sullivan, B., Dinh, K. K., Khuu, N., Hancock, S., Shiu, S.-H., Zipfel, C., Cheung, A. Y., Beilstein, M. A., & Palanivelu, R. (2022). Evolutionary analysis of the *LORELEI* gene family in plants reveals regulatory subfunctionalization. Plant Physiology, 190(4), 2539–2556. 10.1093/plphys/kiac444

Noble, J. A., & Palanivelu, R. (2020). Workflow to Characterize Mutants with Reproductive Defects. In A. Geitmann (Ed.), Pollen and Pollen Tube Biology (Vol. 2160, pp. 109–128). Springer US. 10.1007/978-1-0716-0672-8_8

Panchy, N., Lehti-Shiu, M., & Shiu, S.-H. (2016). Evolution of Gene Duplication in Plants. Plant Physiology, 171(4), 2294–2316. 10.1104/pp.16.00523

Pertea, G., & Pertea, M. (2020). GFF Utilities: GffRead and GffCompare. F1000Research, 9, 304. 10.12688/f1000research.23297.2

Ponvert, N., & Johnson, M. A. (2024). Synergid cell calcium oscillations refine understanding of FERONIA/LORELEI signaling during interspecific hybridization. Plant Reproduction, 37(1), 57–68. 10.1007/s00497-023-00483-6

Ramsey, J., & Schemske, D. W. (1998). PATHWAYS, MECHANISMS, AND RATES OF POLYPLOID FORMATION IN FLOWERING PLANTS. Annual Review of Ecology and Systematics, 29(1), 467–501. 10.1146/annurev.ecolsys.29.1.467

Sandaklie-Nikolova, L., Palanivelu, R., King, E. J., Copenhaver, G. P., Drews, G. N. (2007). Synergid cell death in Arabidopsis is triggered following direct interaction with the pollen tube. Plant Physiol. 144:1753–62. 10.1104/pp.107.098236

Schindelin, J., Arganda-Carreras, I., Frise, E., Kaynig, V., Longair, M., Pietzsch, T., Preibisch, S., Rueden, C., Saalfeld, S., Schmid, B., Tinevez, J.-Y., White, D. J., Hartenstein, V., Eliceiri, K., Tomancak, P., & Cardona, A. (2012). Fiji: An open-source platform for biological-image analysis. Nature Methods, 9(7), 676–682. 10.1038/nmeth.2019

Shen, W., Le, S., Li, Y., & Hu, F. (2016). SeqKit: A Cross-Platform and Ultrafast Toolkit for FASTA/Q File Manipulation. PLOS ONE, 11(10), e0163962. 10.1371/journal.pone.0163962

Shen, W., Sipos, B., & Zhao, L. (2024). SeqKit2: A Swiss army knife for sequence and alignment processing. iMeta, 3(3), e191. 10.1002/imt2.191

Smyth, D. R., Bowman, J. L., & Meyerowitz, E. M. (1990). Early flower development in Arabidopsis. The Plant Cell, 2(8), 755–767. 10.1105/tpc.2.8.755

Suyama, M., Torrents, D., & Bork, P. (2006). PAL2NAL: Robust conversion of protein sequence alignments into the corresponding codon alignments. Nucleic Acids Research, 34(Web Server), W609–W612. 10.1093/nar/gkl315

Takeuchi, H., & Higashiyama, T. (2012). A Species-Specific Cluster of Defensin-Like Genes Encodes Diffusible Pollen Tube Attractants in Arabidopsis. PLoS Biology, 10(12), e1001449. 10.1371/journal.pbio.1001449

Takeuchi, H., & Higashiyama, T. (2016). Tip-localized receptors control pollen tube growth and LURE sensing in Arabidopsis. Nature, 531(7593), 245–248. 10.1038/nature17413

Talavera, G., & Castresana, J. (2007). Improvement of Phylogenies after Removing Divergent and Ambiguously Aligned Blocks from Protein Sequence Alignments. Systematic Biology, 56(4), 564–577. 10.1080/10635150701472164

Tovar-Méndez, A., Lu, L., & McClure, B. (2017). HT proteins contribute to S-RN ase-independent pollen rejection in *Solanum*. The Plant Journal, 89(4), 718–729. 10.1111/tpj.13416

Tsukamoto, T., Qin, Y., Huang, Y., Dunatunga, D., & Palanivelu, R. (2010). A role for LORELEI, a putative glycosylphosphatidylinositol-anchored protein, in Arabidopsis thaliana double fertilization and early seed development: LORELEI functions in fertilization and seed development. The Plant Journal, 62(4), 571–588. 10.1111/j.1365-313X.2010.04177.x

Wang, L., Fan, L., Zhao, Z., Zhang, Z., Jiang, L., Chai, M., & Tian, C. (2022). The Capparis spinosa var. Herbacea genome provides the first genomic instrument for a diversity and evolution study of the Capparaceae family. GigaScience, 11, giac106. 10.1093/gigascience/giac106

Wang, L., & Filatov, D. A. (2023). Mechanisms of prezygotic post-pollination reproductive barriers in plants. Frontiers in Plant Science, 14, 1230278. 10.3389/fpls.2023.1230278

Weadick, C. J., & Chang, B. S. W. (2012). An Improved Likelihood Ratio Test for Detecting Site-Specific Functional Divergence among Clades of Protein-Coding Genes. Molecular Biology and Evolution, 29(5), 1297–1300. 10.1093/molbev/msr311

Williams, E. G., Knox, R. B., & Rouse, J. L. (1982). Pollination sub-systems distinguished by pollen tube arrest after incompatible interspecific crosses in *Rhododendron* (Ericaceae). Journal of Cell Science, 53(1), 255–277. 10.1242/jcs.53.1.255

Wong, W. S. W., Yang, Z., Goldman, N., & Nielsen, R. (2004). Accuracy and Power of Statistical Methods for Detecting Adaptive Evolution in Protein Coding Sequences and for Identifying Positively Selected Sites. Genetics, 168(2), 1041–1051. 10.1534/genetics.104.031153

Xiao, Y., Stegmann, M., Han, Z., DeFalco, T. A., Parys, K., Xu, L., Belkhadir, Y., Zipfel, C., & Chai, J. (2019). Mechanisms of RALF peptide perception by a heterotypic receptor complex. Nature, 572(7768), 270–274. 10.1038/s41586-019-1409-7

Yang, Z. (1998). Likelihood ratio tests for detecting positive selection and application to primate lysozyme evolution. Molecular Biology and Evolution, 15(5), 568–573. 10.1093/oxfordjournals.molbev.a025957

Yang, Z. (2005). Bayes Empirical Bayes Inference of Amino Acid Sites Under Positive Selection. Molecular Biology and Evolution, 22(4), 1107–1118. 10.1093/molbev/msi097

Yang, Z. (2007). PAML 4: Phylogenetic Analysis by Maximum Likelihood. Molecular Biology and Evolution, 24(8), 1586–1591. 10.1093/molbev/msm088

Yang, Z., & Nielsen, R. (1998). Synonymous and nonsynonymous rate variation in nuclear genes of mammals. Journal of Molecular Evolution, 46(4), 409–418. 10.1007/PL00006320

Yang, Z., & Nielsen, R. (2002). Codon-Substitution Models for Detecting Molecular Adaptation at Individual Sites Along Specific Lineages. Molecular Biology and Evolution, 19(6), 908–917. 10.1093/oxfordjournals.molbev.a004148

Yang, Z., Nielsen, R., Goldman, N., & Pedersen, A.-M. K. (2000). Codon-Substitution Models for Heterogeneous Selection Pressure at Amino Acid Sites. Genetics, 155(1), 431–449. 10.1093/genetics/155.1.431

Zhang, J., Nielsen, R., & Yang, Z. (2005). Evaluation of an Improved Branch-Site Likelihood Method for Detecting Positive Selection at the Molecular Level. Molecular Biology and Evolution, 22(12), 2472–2479. 10.1093/molbev/msi237

Zhang, L., Huang, J., Su, S., Wei, X., Yang, L., Zhao, H., Yu, J., Wang, J., Hui, J., Hao, S., Song, S., Cao, Y., Wang, M., Zhang, X., Zhao, Y., Wang, Z., Zeng, W., Wu, H.-M., Yuan, Y., … Duan, Q. (2021). FERONIA receptor kinase-regulated reactive oxygen species mediate self-incompatibility in Brassica rapa. Current Biology, 31(14), 3004–3016.e4. 10.1016/j.cub.2021.04.060

Zhang T., Li S., Dou S., Duan Q. (2026). From recognition to response: integrated signaling pathways determining pollen acceptance and rejection in Brassicaceae. New Phytologist, 250, 788–798. 10.1111/nph.70991

Zhong, S., Li, L., Wang, Z., Ge, Z., Li, Q., Bleckmann, A., Wang, J., Song, Z., Shi, Y., Liu, T., Li, L., Zhou, H., Wang, Y., Zhang, L., Wu, H.-M., Lai, L., Gu, H., Dong, J., Cheung, A. Y., … Qu, L.-J. (2022). RALF peptide signaling controls the polytubey block in *Arabidopsis*. Science, 375(6578), 290–296. 10.1126/science.abl4683

Zhong, S., Liu, M., Wang, Z., Huang, Q., Hou, S., Xu, Y.-C., Ge, Z., Song, Z., Huang, J., Qiu, X., Shi, Y., Xiao, J., Liu, P., Guo, Y.-L., Dong, J., Dresselhaus, T., Gu, H., & Qu, L.-J. (2019). Cysteine-rich peptides promote interspecific genetic isolation in *Arabidopsis*. Science, 364(6443), eaau9564. 10.1126/science.aau9564

