## Supplemental Figures 1-10 for "*LORELEI* sequence variation modulates lineage-dependent pollen tube reception in Brassicaceae"

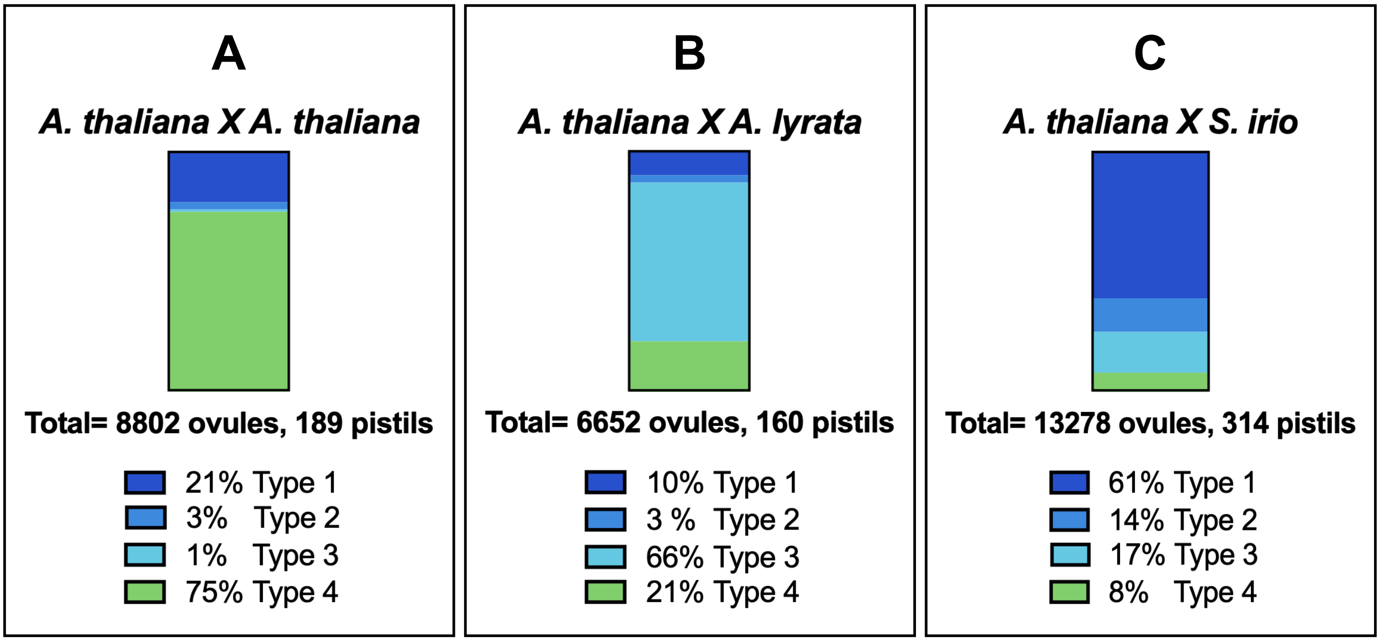


**Figure S1:** Quantification of all types of pollen-pistil interactions during interspecific crosses

(A) *A. thaliana* female was crossed to *A. thaliana* pollen, and four types of pollen-ovule interactions were scored.

(B) *A. thaliana* female was crossed to *A. lyrata* pollen, and four types of pollen-ovule interactions were scored.

(C) *A. thaliana* female was crossed to *S. irio* pollen, and four types of pollen-ovule interactions were scored.

The percentage of each type is listed at the bottom along with the total number of ovules and pistils scored.

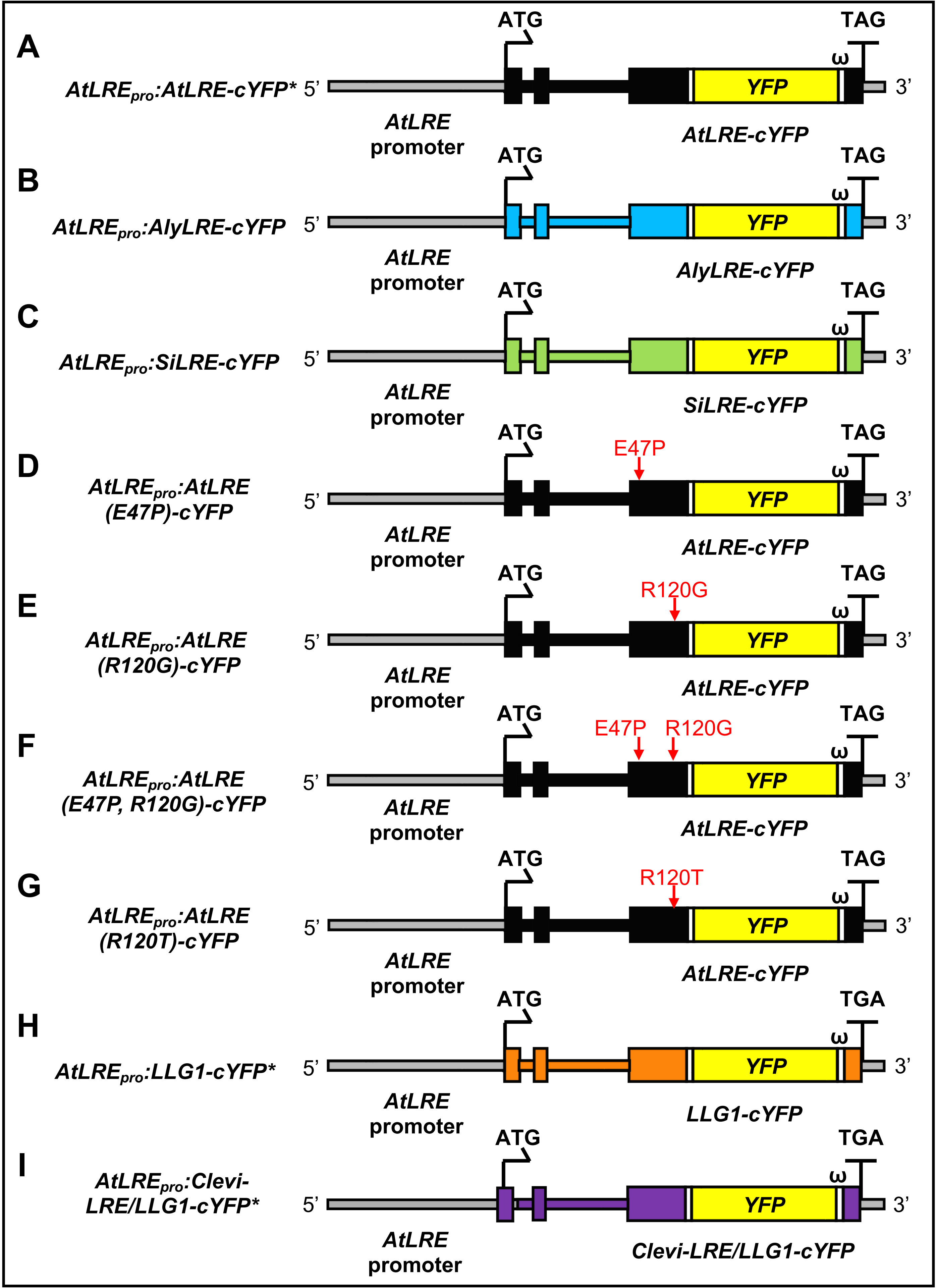

A
AtLREpro:AtLRE-cYFP*
5'
ATG
TAG
ω
YFP
3'
AtLRE promoter
AtLRE-cYFP
B
AtLREpro:AlyLRE-cYFP
AlyLRE-cYFP
C
AtLREpro:SiLRE-cYFP
SiLRE-cYFP
D
AtLREpro:AtLRE (E47P)-cYFP
E47P
E
AtLREpro:AtLRE (R120G)-cYFP
R120G
F
AtLREpro:AtLRE (E47P, R120G)-cYFP
E47P
R120G
G
AtLREpro:AtLRE (R120T)-cYFP
R120T
H
AtLREpro:LLG1-cYFP*
TGA
LLG1-cYFP
I
AtLREpro:Clevi-LRE/LLG1-cYFP*
Clevi-LRE/LLG1-cYFP

**Figure S2:** Constructs introduced into the *A. thaliana lre-7* mutant for generating transgenic plants used in this study.

All constructs used the *A. thaliana* LRE promoter (959 bp), followed by the signal peptide and genomic sequence of the indicated protein. Point mutations in *A. thaliana* LRE (A) were generated using primers (Table S1) that incorporated the desired nucleotide changes (D, E, F, G). Full-length *LRE* sequences from *Sisymbrium irio* and *Arabidopsis lyrata* were obtained from CoGe (https://genomevolution.org/coge/) (B,C). Citrine YFP was fused to the final exon immediately before the predicted omega region and 3' UTR. Previously reported constructs (marked with *) included are $AtLRE_{pro}$*:AtLRE-cYFP* (Liu et al., 2016) and $AtLRE_{pro}$*:LLG1-cYFP* and $AtLRE_{pro}$*:Clevi-LRE/LLG1-cYFP* (Noble et al., 2022).

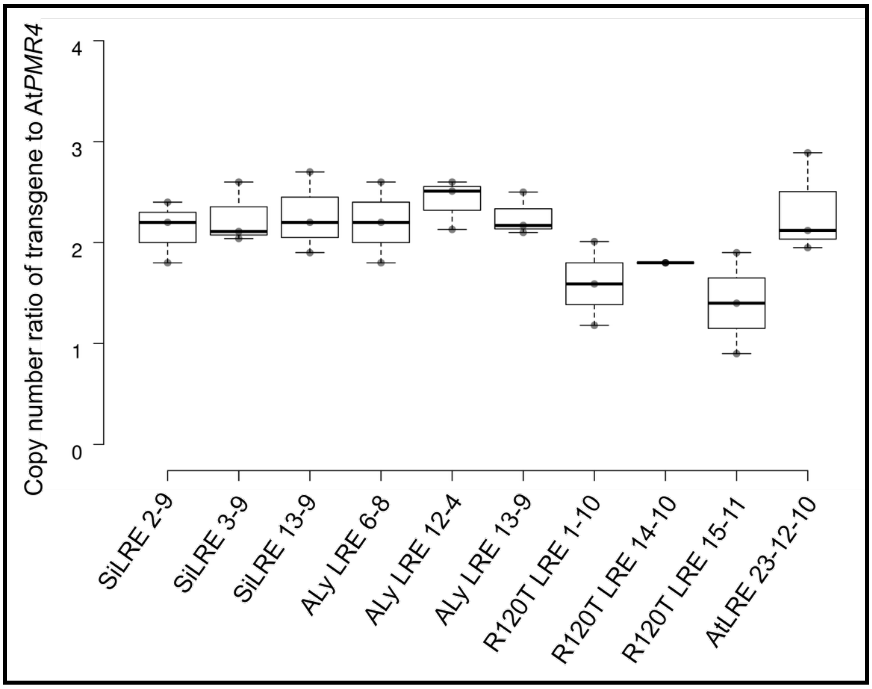


**Figure S3.** ddPCR-based estimation of the number of copies of transgenes present in transgenic lines.

Genomic DNA was extracted from seeds (diploid) of individual lines used in inter-lineage crosses and analyzed for transgene copy number using YFP-specific oligonucleotide probes. The results were compared against reference loci in the Arabidopsis thaliana genome. Box plots represent the copy number of the LRE-YFP transgene in each species, quantified via droplet digital PCR (ddPCR) using the Bio-Rad QX200 system. The endogenous reference gene AtPMR4 (AT4G03550), a diploid gene with a known copy number of two, was used for normalization. A copy number ratio close to 2.0 suggests the presence of two copies at a single genomic locus (co-segregating insertion). Error bars represent standard deviation (SD) across three biological replicates. The center lines of the box plots show the medians, and box edges represent the 25th and 75th percentiles.

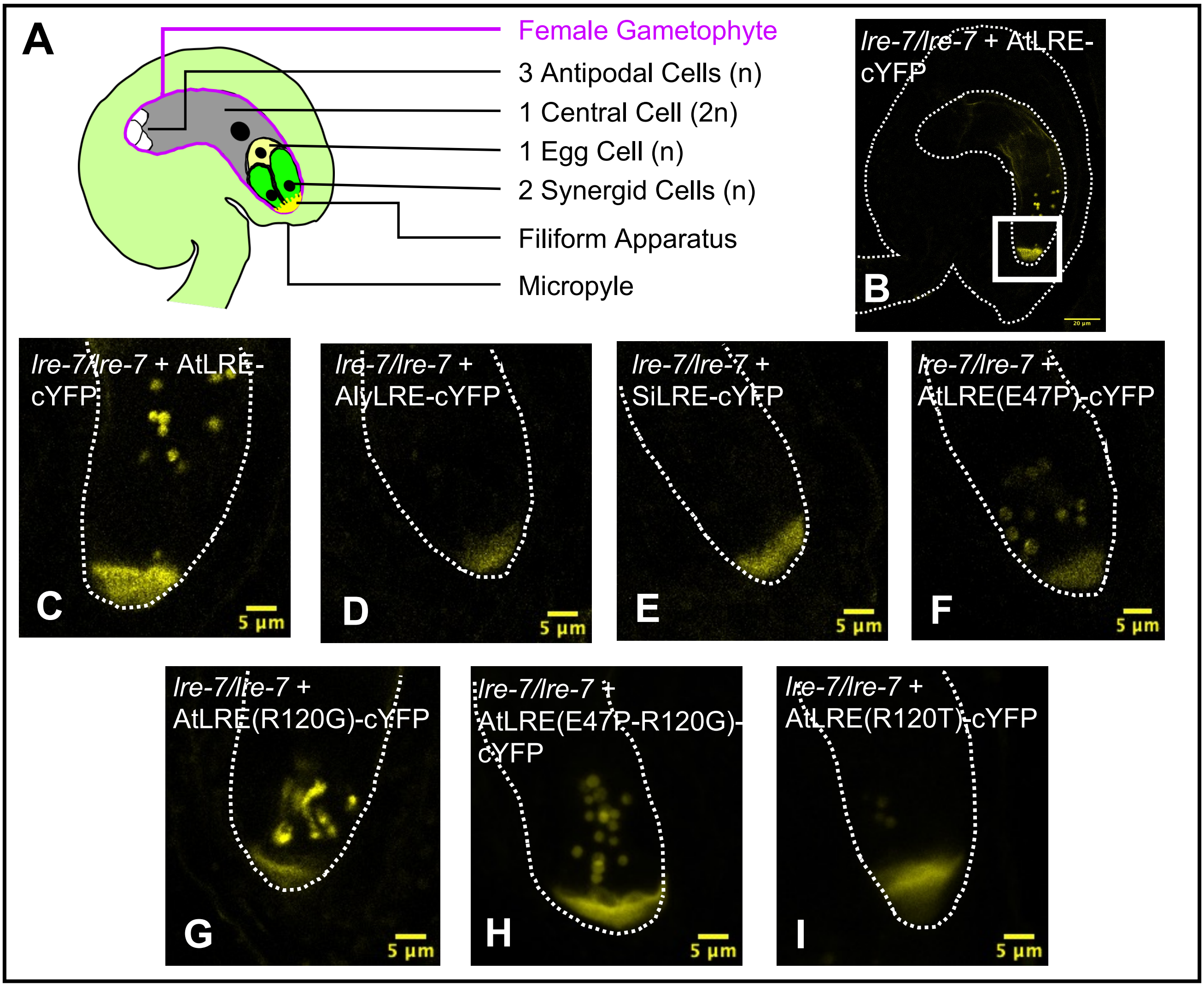


**Figure S4: LRE-cYFP expressed in the transgenic plants localize to the filiform apparatus of the synergid cells in the ovules of unpollinated pistils.**

(A) Diagram of a mature ovule before pollination showing it encasing a mature female gametophyte, composed of seven cells belonging to four distinct types: three haploid antipodal cells, one large diploid central cell, one haploid egg cell, and two haploid synergid cells, which share a highly membranous and secretory structure, the filiform apparatus. The micropyle is the opening in the ovule through which the pollen tube will enter to deliver the sperm cells and complete fertilization.

(B, C) As previously reported, a confocal image of an ovule carrying the *pLRE::AtLRE-cYFP* construct shows LRE-cYFP expressed from it localizes to the filiform apparatus of the synergid cells (Liu et al., 2016). White rectangle highlights the area shown in C and similar micropylar regions of corresponding ovules in D-I.

(D-H) Confocal images showing expression of citrine YFP signals from six other constructs used in this study also localizing to the filiform apparatus: *AtLRE$_{pro}$:AlyLRE-cYFP* (D), *AtLRE$_{pro}$ :SiLRE-cYFP* (E), *AtLRE$_{pro}$ :AtLRE (E47P)-cYFP* (F), *AtLRE$_{pro}$ :AtLRE (R120G)-cYFP* (G), *AtLRE$_{pro}$ :AtLRE (E47P, R120G)-cYFP* (H), and *AtLRE$_{pro}$ :AtLRE (R120T)-cYFP* (I).

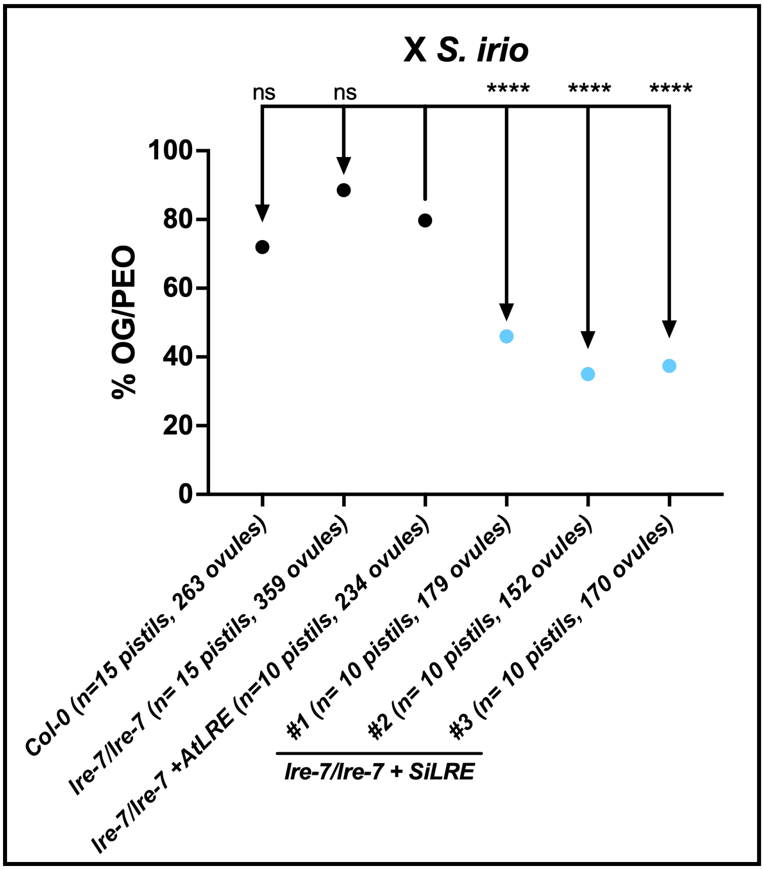


**Figure S5: *A. thaliana lre-7* mutant ovules complemented with full length LRE from *S. irio* significantly rescued pollen tube overgrowth phenotype when crossed to *S. irio***

Quantification of the average pollen tube overgrowth per silique (%OG/PEO) between *AtLRE$_{pro}$:SiLRE-cYFP* female parents and S. irio pollen. The average % OG/PEO for each female was compared to % OG/PEO of *lre-7/lre-7+ AtLRE$_{pro}$:AtLRE-cYFP*. In this construct, three independently transformed lines were crossed to *S. irio*. % OG/PEO indicates the percentage of ovules in a silique that experienced pollen tube overgrowth in the female gametophyte out of all the ovules that received a pollen tube. The number of pistils crossed, and ovules scored is listed along with each female parent on the X-axis. Statistical significance was determined by first using an arcsine transformation of percentage data, followed by pairwise t-tests between *lre-7/lre-7+ AtLRE$_{pro}$:AtLRE-cYFP* and tested lines; ns> 0.05, *p<0.05, **p<0.01, ***p<0.001, and ****p<0.0001. This experiment was conducted at the Czech Academy of Sciences in the Institute of Pollen Biology.

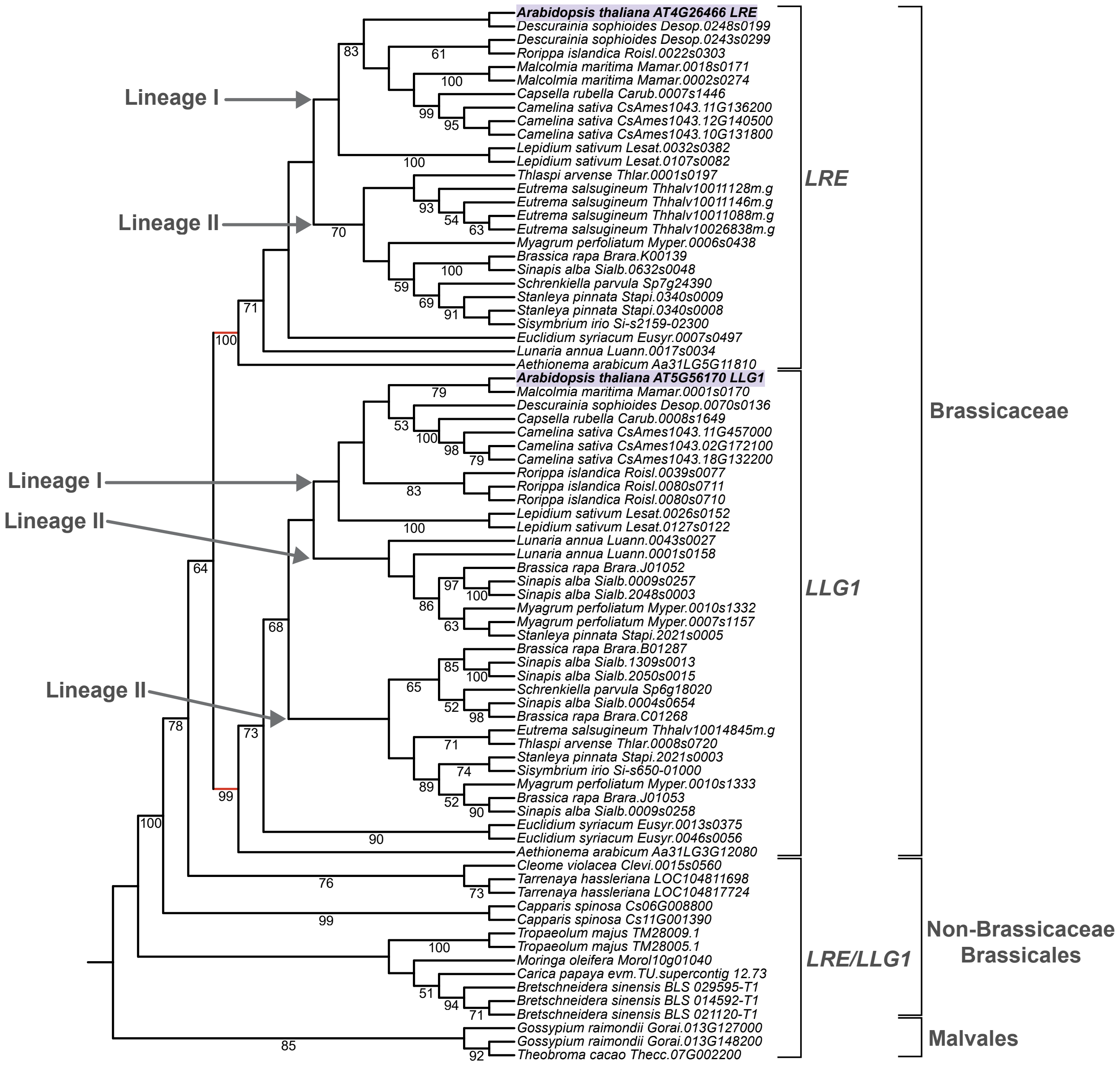


**Figure S6: Cladogram with bootstrap support of *LRE* and *LLG1* homologues in Brassicales and Malvales.**

Maximum-likelihood cladogram inferred from 80 *LRE* and *LLG1* homologues retrieved from a set of 27 Brassicales + Malvales species. The tree was rooted on Malvales *LRE/LLG1* homologues. *LRE* and *LLG1* clades from Lineage I and Lineage II Brassicaceae are marked with grey arrows. *LRE* and *LLG1* post-duplication branches are marked in red. *LRE* and *LLG1* from *A. thaliana* are highlighted with purple. Node support was determined using 1000 Felsenstein bootstrap replicates (bootstrap values over 50% are reported).

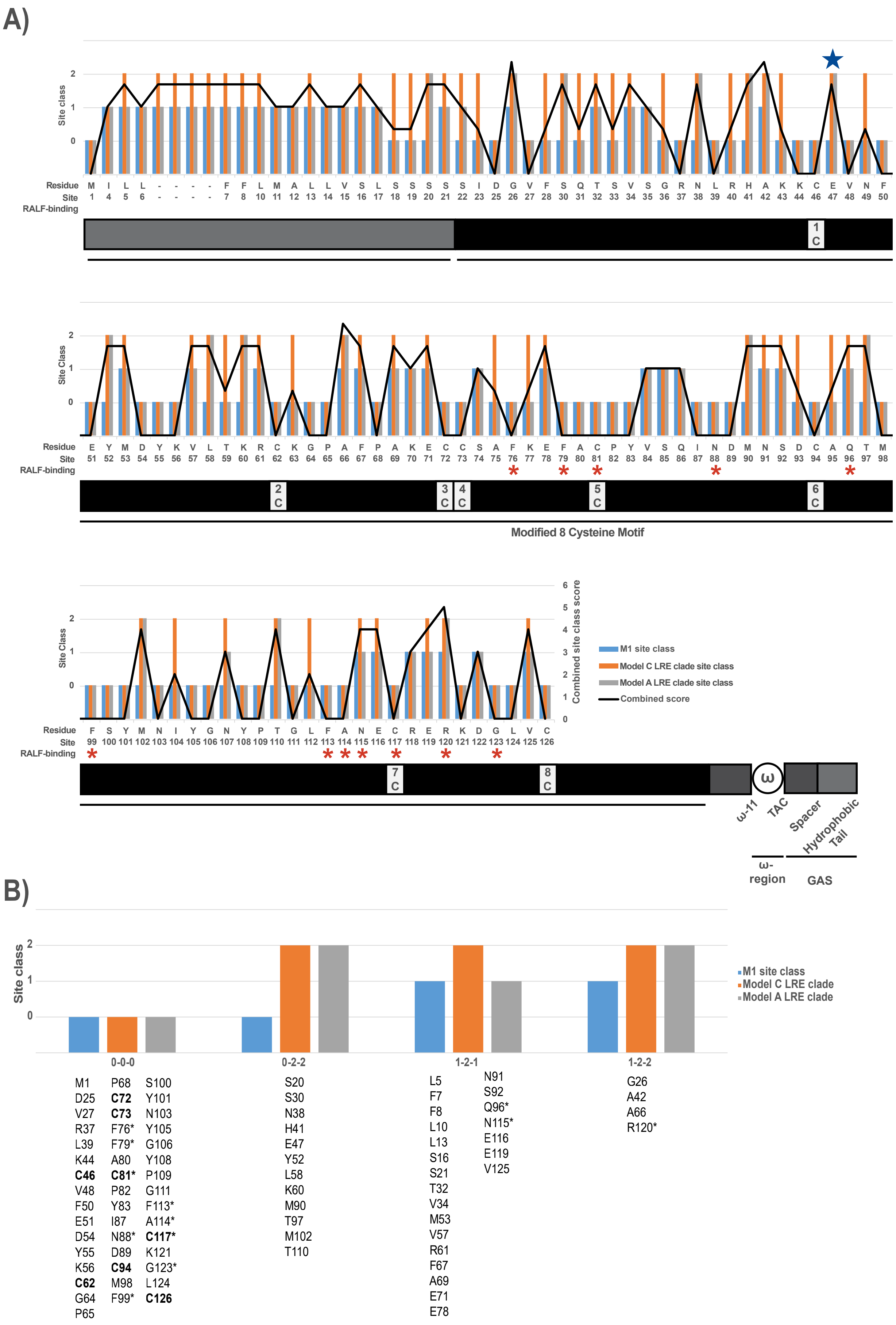
A)
Site class
Residue
Site
RALF-binding
Site Class
Modified 8 Cysteine Motif
Combined site class score
M1 site class
Model C LRE clade site class
Model A LRE clade site class
Combined score
ω-11
TAC
Spacer
Hydrophobic Tail
ω-region
GAS
B)
Site class
M1 site class
Model C LRE clade
Model A LRE clade
0-0-0
M1 D25 V27 R37 L39 K44 C46 V48 F50 E51 D54 Y55 K56 C62 G64 P65
P68 C72 C73 F76* F79* A80 C81* P82 Y83 I87 N88* D89 C94 M98 F99*
S100 Y101 N103 Y105 G106 Y108 P109 G111 F113* A114* C117* K121 G123* L124 C126
0-2-2
S20 S30 N38 H41 E47 Y52 L58 K60 M90 T97 M102 T110
1-2-1
L5 F7 F8 L10 L13 S16 S21 T32 V34 M53 V57 R61 F67 A69 E71 E78
N91 S92 Q96* N115* E116 E119 V125
1-2-2
G26 A42 A66 R120*

**Figure S7** Selective pressures on certain sites changed in *LRE* post-duplication.

(A) Results of Bayes Empirical Bayes analyses are shown as a function of each residue's position, amino acid, and domain in At*LRE*. Site classes are as follows: for the site model M1 (Blue bars), site class 0 corresponds to $\omega_0$, and site class 1 to $\omega_1$; for the clade Model C with the *LRE* clade as foreground (Orange bars), site class 0 corresponds to $\omega_0$, site class 1 to $\omega_1$, and site class 2 to $\omega_{2\_background}$ and $\omega_{2\_LRE}$; for the branch-sites Model A with the *LRE* clade as foreground (Grey bars), site class 0 corresponds to $\omega_0$, site class 1 to $\omega_1$, site class 2 to $\omega_{2a\_background}$ and $\omega_{2a\_LRE}$, and site class 3 to $\omega_{2b\_background}$ and $\omega_{2b\_LRE}$. A combined site class score (Black line) for each site was determined by adding up the value of the three previously mentioned BEB scores. Values for the site classes can be found in Table S5. Cysteines in the Modified 8 Cysteine Motif (M8CM) are numbered, and sites identified by Xiao et al., 2019 to facilitate LLG1/2/3 binding with RALF23 are noted with asterisks. A blue star marks E47, which is the only site that belongs to site class 2 when the branch-sites Model A test was run with the *LRE* PDB as the foreground.

(B) Four distinct site class profiles, based on BEB analyses in (A). Site class profiles 0-0-0, 0-2-2, 1-2-1, and 1-2-2 each represent a given residue's site class, where the first number corresponds to site class for M1, and where the second and third numbers correspond to the site class for Model C with with *LRE* clade as foreground and Model A with *LRE* clade as foreground, respectively. Bolded residues represent cysteines that are a part of the M8CM, and residues marked with an asterisk were identified by Xiao et al., 2019 to facilitate LLG1/2/3 binding with RALF23.

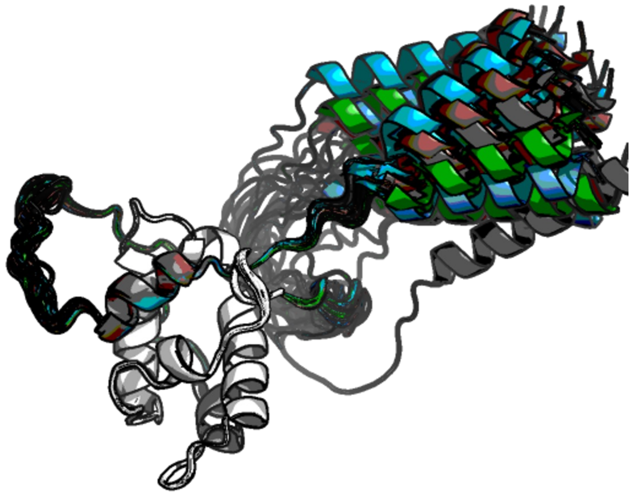

**Figure S8:** The AlphaFold3-predicted ground state structures of WT *At*LRE and variants show little-to-no variation between structures.

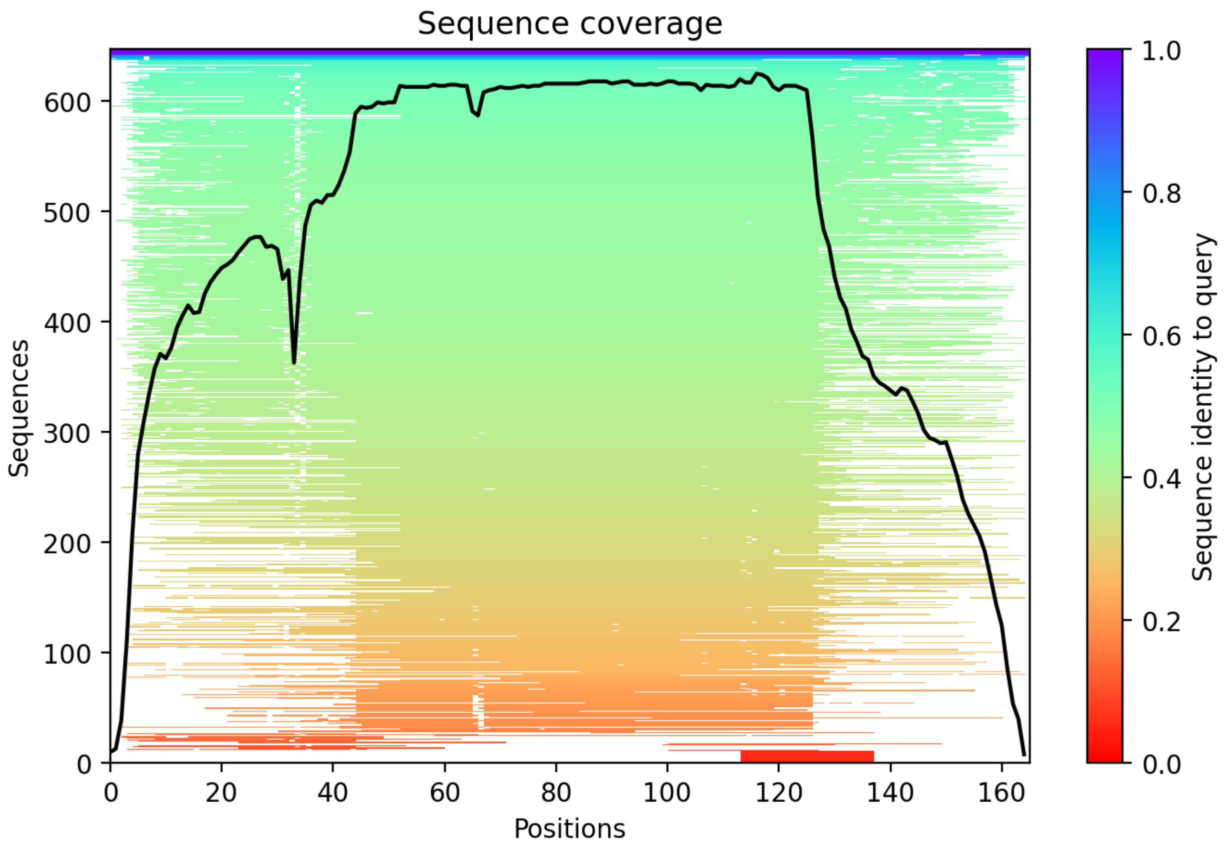


**Figure S9:** Multi-sequence alignment coverage of WT *At*LRE shows low sequence coverage and identity at the N- and C-termini.

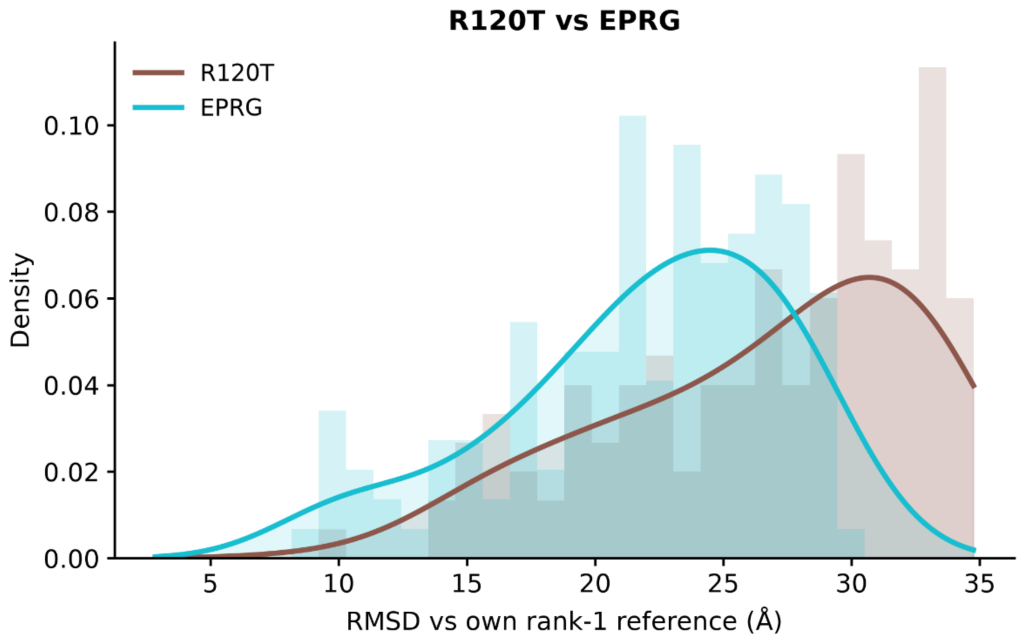


**Figure S10:** Overlay of 1D RMSD of EGRP and R120T, each calculated in relation to its own highest-ranked structures.
