## Supplemental Table S3 for "*LORELEI* sequence variation modulates lineage-dependent pollen tube reception in Brassicaceae"

| **Table S3.** Post gene duplication, the *LRE* clade, but not the *LLG1* clade, experienced a relaxation in purifying selection | | | | | |
|---|---|---|---|---|---|
| Test | Branch(es) highlighted[a] | Alternative hypothesis (lnL) | Null hypothesis (lnL) | LRT Score[b] | *p*-value |
| Sites test (M1) | | | | | |
| | NA | -10098.09995 | -10229.96359 | 263.727284 | < 0.0001** |
| Branch test | | | | | |
| | *LRE* PDB | -10229.54446 | -10229.96359 | 0.838258 | 0.3599 |
| | *LRE* Clade | -10226.43249 | -10229.96359 | 7.062192 | 0.0079* |
| | *LLG1* PDB | -10229.46662 | -10229.96359 | 0.993934 | 0.3188 |
| | *LLG1* Clade | -10229.58676 | -10229.96359 | 0.753662 | 0.3853 |
| Model C clade test | | | | | |
| | *LRE* Clade | -10024.14561 | -10027.69313 | 7.095034 | 0.0077* |
| Model A branch-sites test | | | | | |
| | *LRE* PDB | -10095.58497 | -10097.07992 | 2.989912 | 0.0838 |
| | *LRE* Clade | -10064.0625 | -10064.0625 | 0 | 1 |

[a] *LRE* and *LLG1* clades and post-duplication branches (PDBs) are indicated in Figure 1b
[b] Likelihood Ratio Test (LRT) score was calculated by taking difference squared of the likelihood ratio (lnL) scores of the alternative hypothesis lnL compared to that of the null hypothesis lnL
[c] LRT p-values are determined using a Chi-square test (for all tests performed, DF=1)
*$p$<0.05, **$p$<0.005
