## Supplemental Table S4 for "*LORELEI* sequence variation modulates lineage-dependent pollen tube reception in Brassicaceae"

| **Table S4.** Bayes Empirical Bayes results of sites under relaxed selection for Model A tests performed on *LRE* clade branches and their relative positions in *LRE* and *LLG1* | | | | |
|---|---|---|---|---|
| Amino acid residue in the alignment | BEB score with *LRE* PBD as foreground[a] | BEB score with *LRE* clade as foreground[a] | Position relative to *A. thaliana LRE* | Position relative to *A. thaliana LLG1* |
| 21 S | 0.977 | NS | S20 | S22 |
| 26 G | 0.646 | NS | G26 | G29 |
| 29 S | 0.921 | NS | S30 | S33 |
| 37 N | 0.822 | NS | N38 | N41 |
| 40 H | 0.949 | NS | H41 | Q44 |
| 41 A | 0.581 | NS | A42 | T45 |
| 45 E | 0.923 | 0.953 | E47[b] | P50 |
| 50 Y | 0.932 | NS | Y52 | F55 |
| 56 L | 0.990 | NS | L58 | I61 |
| 58 K | 0.796 | NS | K60 | S63 |
| 64 A | 0.640 | NS | A66 | K69 |
| 88 M | 0.917 | NS | M90 | L93 |
| 95 T | 0.937 | NS | T97 | T100 |
| 100 M | 0.999 | NS | M102 | I105 |
| 108 T | 0.998 | NS | T110 | P113 |
| 118 R | 0.618 | NS | R120[b] | G123 |

[a]Bayes Empirical Bayes (BEB) analyses identified positions in the translation alignment that had an increased evolutionary rate in the highlighted branch(es) for the Model A branch-site test (Yang et al., 2005).

[b]E47 and R120 were selected for further analysis.

NS, no score generated by BEB analysis.
