## Supplemental Table S6 for "*LORELEI* sequence variation modulates lineage-dependent pollen tube reception in Brassicaceae"

**Table S6.** List of primers used in this study

| Primer | Primer Name | Sequence (5' - 3') | Template | Primer Pair | Expected Length (bp) |
|---|---|---|---|---|---|
| ***AtLRE*$_{pro}$*:AthLRE(E47P)-cYFP* construct** | | | | | |
| 1701 | *AtLRE*$_{pro}$*:LRE-cYFP* FWD Primer 1 | AGGCGGCCGCACTAGTATCTGTGAGTCATCCTTTCGAGGAAATC | *AtLRE*$_{pro}$*:LRE-cYFP* plasmid | 1701 + 1966 | 1615 |
| 1966 | *AtLRE*$_{pro}$*:LRE-E47P-R120G-cYFP* REV Primer 1 | CTCAAAGTTCACAGGACATTCTGTTTCCAACCAAACA | | | |
| 1967 | *AtLRE*$_{pro}$*:LRE-E47P-cYFP* FWD Primer 2 | CCTGTGAACTTTGAGTATATGGACTACAAGGT | *AtLRE*$_{pro}$*:LRE-cYFP* plasmid | 1967 + 1702 | 1288 |
| 1702 | *AtLRE*$_{pro}$*:LRE-cYFP* REV Primer 2 | AGCTGGGTCGGCGCGCCGGAGGTCAAGTATTCTTTACACTTGGACACT | | | |
| ***AtLRE*$_{pro}$*:AthLRE(R120T)-cYFP* construct** | | | | | |
| 1701 | *AtLRE*$_{pro}$*:LRE-cYFP* FWD Primer 1 | AGGCGGCCGCACTAGTATCTGTGAGTCATCCTTTCGAGGAAATC | *AtLRE*$_{pro}$*:LRE-cYFP* plasmid | 1701 + 1972 | 1835 |
| 1972 | *AtLRE*$_{pro}$*:LRE-R120T-cYFP* REV Primer 1 | CAAGCCCATCTTTTGTTTCTCTGCACTCGTTAGC | | | |
| 1973 | *AtLRE*$_{pro}$*:LRE-R120T-cYFP* FWD Primer 2 | ACAAAAGATGGGCTTGTTTGCCCT | *AtLRE*$_{pro}$*:LRE-cYFP* plasmid | 1973 + 1702 | 1069 |
| 1702 | *AtLRE*$_{pro}$*:LRE-cYFP* REV Primer 2 | AGCTGGGTCGGCGCGCCGGAGGTCAAGTATTCTTTACACTTGGACACT | | | |
| ***AtLRE*$_{pro}$*:AthLRE(R120G)-cYFP* construct** | | | | | |
| 1701 | *AtLRE*$_{pro}$*:LRE-cYFP* FWD Primer 1 | AGGCGGCCGCACTAGTATCTGTGAGTCATCCTTTCGAGGAAATC | *AtLRE*$_{pro}$*:LRE-cYFP* plasmid | 1701 + 1968 | 1835 |
| 1968 | *AtLRE*$_{pro}$*:LRE-E47P-R120G-REV Primer 2* | CAAGCCCATCTTTACCTTCTCTGCACTCGTTAGCG | | | |
| 1969 | *AtLRE*$_{pro}$*:LRE-E47P-R120G-FWD Primer 3* | GGTAAAGATGGGCTTGTTTGCCCTTTA | *AtLRE*$_{pro}$*:LRE-cYFP* plasmid | 1969 + 1702 | 1069 |
| 1702 | *AtLRE*$_{pro}$*:LRE-cYFP* REV Primer 2 | AGCTGGGTCGGCGCGCCGGAGGTCAAGTATTCTTTACACTTGGACACT | | | |
| ***AtLRE*$_{pro}$*:AthLRE(E47P-R120G)-cYFP* construct** | | | | | |
| 1701 | *AtLRE*$_{pro}$*:LRE-cYFP* FWD Primer 1 | AGGCGGCCGCACTAGTATCTGTGAGTCATCCTTTCGAGGAAATC | *AtLRE*$_{pro}$*:LRE-cYFP* plasmid | 1701 + 1966 (PCR | 1615 |

| | | | | | |
|---|---|---|---|---|---|
| 1966 | *AtLRE$_{pro}$:LRE-E47P-R120G-REV Primer 1* | CTCAAAGTTCACAGGACATTCTGTTTCCAACCAAACA | | Product 1) | |
| 1967 | *AtLRE$_{pro}$:LRE-E47P-R120G-FWD Primer 2* | CCTGTGAACTTTGAGTATATGGACTACAAGGT | *AtLRE$_{pro}$:LRE-cYFP* plasmid | 1967 + 1968 (PCR Product 2) | 235 |
| 1968 | *AtLRE$_{pro}$:LRE-E47P-R120G-REV Primer 2* | CAAGCCCATCTTTACCTTCTCTGCACTCGTTAGCG | | | |
| 1969 | *AtLRE$_{pro}$:LRE-E47P-R120G-FWD Primer 3* | GGTAAAGATGGGCTTGTTTGCCCTTTA | *AtLRE$_{pro}$:LRE-cYFP* plasmid | 1969 + 1702 (PCR Product 3) | 1069 |
| 1702 | *AtLRE$_{pro}$:LRE-cYFP* REV Primer 2 | AGCTGGGTCGGCGCGCCGGAGGTCAAGTATTCTTTACACTTGGACACT | | | |
| *Overlap PCR Fragments 2+3* | | | PCR products 2 + 3 | 1967 + 1702 | 1288 |
| ***AtLRE$_{pro}$:AlyLRE-cYFP* construct** | | | | | |
| 1852 | *AtLRE$_{pro}$:AlyLRE-FWD Primer 1* | GCAGGCGGCCGCACTAGTCACAATTCCATGATCTGTGAGTC | *Col-0* genomic DNA | 1852 + 1853 (PCR Product 1) | 1000 |
| 1853 | *AtLRE$_{pro}$:AlyLRE-REV Primer 1* | ATCAGCTCCATGAAATTGTTGTTAAAGAAGCTTGTAACAT | | | |
| 1854 | *AtLRE$_{pro}$:AlyLRE-FWD Primer 2* | ACAACAATTTCATGGAGCTGATATTATTATTCTTCCTTCT | *A. lyrata* genomic DNA | 1854 + 1863 (PCR Product 2) | 875 |
| 1863 | *AlyLRE_Intron_2REV* | ACAAAAGTTATCTCGTAATTACTCTCGG | | | |
| 1862 | *AlyLRE_Intron_2FWD* | CCGAGAGTAATTACGAGATAACTTTTGT | *A. lyrata* genomic DNA | 1862 + 1855 (PCR Product 3) | 1171 |
| 1855 | *AtLRE$_{pro}$:AlyLRE-REV Primer 2* | GATGTGGGGAGTATAGTGGAGGTAGAGGGCAAACAATTCCATCTTTC | | | |
| 1856 | *AtLRE$_{pro}$:AlyLRE-FWD Primer 3* | CCTCCACTATACTCCCCACATCTCAACGCGGGCCGGCCTGGAGGT | *AtLRE$_{pro}$:LRE-cYFP* plasmid | 1856 + 1857 (PCR Product 4) | 785 |
| 1857 | *AtLRE$_{pro}$:AlyLRE-REV Primer 3* | TAGAGAGGTGGTAACTTGTACAGCTCGTCCATGC | | | |
| 1858 | *AtLRE$_{pro}$:AlyLRE-FWD Primer 4* | GTACAAGTTACCACCTCTCTATTCTCCTCACC | *A. lyrata* genomic DNA | 1858 + 1859 (PCR Product 5) | 137 |
| 1859 | *AtLRE$_{pro}$:AlyLRE-REV Primer 4* | CTTTGATTGATCAAGTCAACACTAATAAAGCCAAAACA | | | |
| 1860 | *AtLRE$_{pro}$:AlyLRE-FWD Primer 5* | GTTGACTTGATCAATCAAAGGAAATTGAAAGAGCC | *Col-0* genomic DNA | 1860 + 1861 (PCR Product 6) | 301 |
| 1861 | *AtLRE$_{pro}$:AlyLRE-REV Primer 5* | AAGCTGGGTCGGCGCGCCTGTACCCACATGACTTTTGATTTTTGATG | | | |

| | | | | | |
|---|---|---|---|---|---|
| *Overlap PCR Fragments 1+2* | | | PCR products 1+2 | 1852 + 1863 | 1853 |
| *Overlap PCR Fragments 5+6* | | | PCR products 5+6 | 1858 + 1861 | 418 |
| *Overlap PCR Fragments 4+5+6* | | | PCR products 4+(5+6) | 1856 + 1861 | 1182 |
| *Overlap PCR Fragments 3+4+5+6* | | | PCR products 3+(4+5+6) | 1862 + 1861 | 2331 |
| ***$AtLRE_{pro}$:SirLRE-cYFP* construct** | | | | | |
| 1701 | *$AtLRE_{pro}$:LRE-cYFP* FWD Primer 1 | AGGCGGCCGCACTAGTATCTGTGAGTCATCCTTTCGAGGAAATC | *$AtLRE_{pro}$:LRE-cYFP* plasmid | 1701 + 2318 (PCR Product 1) | 990 |
| 2318 | *$AtLRE_{pro}$:SirLRE-cYFP REV Primer 1* | CAATTTTACCTCCATGAAATTGTTGTTAAAGAAGCTTGTAAC | | | |
| 2319 | *$AtLRE_{pro}$:SirLRE-cYFP FWD Primer 2* | ATGGAGGTAAAATTGTTATTCCTTCTGATGGC | *S. irio* genomic DNA | 2319 + 2320 (PCR Product 2) | 1094 |
| 2320 | *$AtLRE_{pro}$:SirLRE-cYFP REV Primer 2* | ACCTCCAGGCCGGCCGCCGTTTAGGTTTGGTG | | | |
| 2321 | *$AtLRE_{pro}$:SirLRE-cYFP FWD Primer 3* | GGCCGGCCTGGAGGT | *$AtLRE_{pro}$:LRE-cYFP* plasmid | 2321 + 2322 (PCR Product 3) | 780 |
| 2322 | *$AtLRE_{pro}$:SirLRE-cYFP REV Primer 3* | ACCATTGAGATTAGGAGATCCAAGTGGAGGTGACGATCGCTTGTACAGCTC | | | |
| 2323 | *$AtLRE_{pro}$:SirLRE-cYFP FWD Primer 4* | CGATCGTCACCTCCACTTGGATCTCCTAATCTCAATGGTTCAACCGCTGATTCGAC | *S. irio* genomic DNA | 2323 + 2324 (PCR Product 4) | 141 |
| 2324 | *$AtLRE_{pro}$:SirLRE-cYFP REV Primer 4* | ATTTCCTTTGATTGACTAAGATAGCAATAGAAAAGCCAA | | | |
| 2325 | *$AtLRE_{pro}$:SirLRE-cYFP FWD Primer 5* | TCAATCAAAGGAAATTGAAAGAGCC | *$AtLRE_{pro}$:LRE-cYFP* plasmid | 2325 + 1702 (PCR Product 5) | 148 |
| 1702 | *$AtLRE_{pro}$:LRE-cYFP* REV Primer 2 | AGCTGGGTCGGCGCGCCGGAGGTCAAGTATTCTTTACACTTGGACACT | | | |
| *Overlap PCR Fragments 1 + 2* | | | PCR products 1 + 2 | 1701 + 2320 | 2069 |
| *Overlap PCR Fragments 4+5* | | | PCR products 4 + 5 | 2323 + 1702 | 274 |

| | | |
|---|---|---|
| *Overlap PCR Fragments 3+4+5* | PCR products 3+(4+5) | 2321 + 1015 1702 |
